# Ancestral origin and structural characteristics of non-syntenic homologous chromosomes in abalones

**DOI:** 10.1101/2025.04.02.641744

**Authors:** Shotaro Hirase, Takashi Makino, Takeshi Takeuchi, Mitsutaka Kadota, Shigehiro Kuraku, Kiyoshi Kikuchi

## Abstract

Structural variation is increasingly recognized as a pivotal contributor to genomic diversity in marine invertebrates, yet its extent and evolutionary significance remain poorly characterized in many species. Haplotype-phased genome assembly is an excellent method for studying such variations by comparing homologous chromosomes. In this study, we constructed a haplotype-phased genome assembly for the western Pacific abalone, *Haliotis gigantea*, using high-fidelity (HiFi) long-read sequencing and high-resolution chromosome conformation capture (Hi-C) data. The primary and alternative assemblies each comprised 18 long scaffolds (>50 Mb), consistent with the species’ diploid chromosome number (2*n* = 36), and contained 96.5% and 96.2% complete single-copy Metazoa Benchmarking Universal Single-Copy Orthologs genes, respectively, indicating high assembly quality. Comparative analysis of the two haplotypes revealed three homologous chromosomes with large-scale non-syntenic regions caused by extensive segmental duplications, with each enriched in distinct gene domains. These non-syntenic chromosomes likely originated in abalone evolution, as they were conserved across both closely and distantly related species. Our findings highlight the evolutionary importance of non-syntenic structural variation in shaping genome architecture and suggest that such variation may play a broader role in functional diversification across abalones.

## Introduction

Marine invertebrates represent a significant portion of global biodiversity (Snelgrove 1999; Eisenhauer and Hines 2021; Chen 2021; GIGA Community of Scientist 2014), and their external morphologies (Lv et al. 2019) and ecological features (Sanford and Kelly 2011) are highly diverse. As if to reflect this diversity, early allozyme studies suggested that marine invertebrate genomes generally exhibit high heterozygosity (Solé-Cava and Thorpe 1991), a finding that has been supported by recent genomic studies (Gerdol et al. 2020; Rosa et al 2015). Furthermore, long-read sequencing has revealed numerous large-scale structural variations, such as inversions, in marine invertebrate genomes (Koch et al. 2021). High-quality genome assemblies are therefore essential to elucidate the genomic features underlying this diversity. Advances in genome sequencing technologies have led to a rapid accumulation of genomic information for various marine invertebrate species (Lopez et al. 2019).

Gene duplication increasingly contributes significantly to genomic diversity in marine invertebrates (Corrochano-Fraile et al. 2022; Makino and Kawata 2019; Takeuchi et al. 2022; Qi et al. 2021; Hirase et al. 2023; Marlétaz et al. 2023; Farhat et al. 2022; Yoshida et al. 2011; Kenny et al. 2016; Pardos-Blas et al. 2021). Gene duplication is a major evolutionary mechanism, facilitating the emergence of novel gene functions and increased gene expression through release from functional constraints (Ohno 1970). Such duplication can occur through whole-genome duplication (WGD) or small-scale duplication (SSD) (Satake et al. 2012). Although WGD events are relatively rare in marine invertebrates (Kenny et al. 2016; Pardos-Blas et al. 2021), recent studies have increasingly identified SSDs in various taxa (Corrochano-Fraile et al. 2022; Makino and Kawata 2019; Takeuchi et al. 2022; Qi et al. 2021; Hirase et al. 2023; Marlétaz et al. 2023; Farhat et al. 2022; Yoshida et al. 2011). These findings suggest that taxon- or species-specific SSD events may have occurred in marine invertebrates, potentially influencing their evolutionary trajectories through the proliferation of duplicate genes. Gene copy number variation, a common outcome of such duplications, has been implicated in environmental adaptation (Dorant et al. 2020), disease resistance (Takeuchi et al. 2022), and speciation (Hirase et al. 2023).

*Haliotis*, commonly referred to as “abalone,” is a genus of marine herbivorous gastropods found in tropical and temperate coastal waters across all continents except the Pacific coast of South America and the Atlantic coast of North America. Approximately 60 species have been described (Geiger 1999). Abalones hold significant ecological, historical, and cultural value and are commercially important in many regions (Geiger and Owen 2012). However, in recent decades, wild abalone populations have undergone drastic declines due to illegal fishing, pollution, climate change, and disease outbreaks (Cook 2019), placing many species at risk of extinction (IUCN, 2022). In response, genome assemblies for various abalone species have been reported to support conservation and population management efforts (Nam et al. 2017; Orland et al. 2022; Griffiths et al. 2022; Botwright et al. 2019; Tshilate et al. 2023; Barkan et al. 2024; Gan et al. 2019; Masonbrink et al. 2019).

Previous studies have suggested that gene duplications may be particularly prevalent in western Pacific abalone species (Hirase et al. 2023), with potential implications for disease resistance and resilience to environmental stressors (Makino and Kawata 2012; Takeuchi et al. 2022). To address the limited understanding of structural variation and gene duplication in abalone genomes, we constructed a haplotype-phased genome assembly of the western Pacific abalone *Haliotis gigantea*. Haplotype-phased genome assembly allow for direct comparison between pairs of homologous chromosomes and are particularly useful for detecting structural variations such as non-syntenic regions (Takeuchi et al. 2022). In this study, we identified three homologous chromosomes with large-scale non-syntenic regions, which appear to be conserved across multiple abalone species and likely originated through segmental duplications. These findings provide new insights into the genomic architecture and evolutionary history of abalone species.

## Materials and Methods

### Genome sequencing and assembly

A wild *H. gigantea* male collected from the coastal area of Awaji Island, Hyogo Prefecture, Japan, was used in this study. Foot muscle tissue was sampled, immediately frozen, and stored at −80°C until DNA extraction. High-molecular-weight genomic DNA was extracted using the phenol/chloroform method with TNES-urea buffer (Asahida et al. 1996) and subsequently purified using the DNeasy PowerClean Pro kit (Qiagen, Hilden, Germany). Short DNA fragments were removed using Circulomics Short Read Eliminator kit (Circulomics, MD, USA). To evaluate DNA quality, the concentration of extracted genomic DNA was measured using a Qubit Fluorometer (Thermo Fisher Scientific, Waltham, MA, USA), and the size distribution of DNA fragments was assessed using a TapeStation 2100 system (Agilent Technologies, Wilmington, DE, USA). High-fidelity (HiFi) SMRTbell libraries were constructed and single-molecule sequencing in circular consensus sequencing (CCS) mode of PacBio Sequel II were performed at BGI (Hong Kong, China). The HiFi reads were assembled using hifiasm ver. 0.16.1 with default parameters. Following Takeuchi et al. (2022), the primary unitigs file (*. p_utg.gfa), which contain the nucleotide sequences of haplotype pairs generated by hifiasm, were used as contigs for subsequent scaffolding with high-resolution chromosome conformation capture (Hi-C) data.

### High-resolution chromosome conformation capture data production and genome scaffolding

A Hi-C library was prepared using foot muscle tissue, following the iconHi-C protocol and employing the restriction enzymes DpnII and HinfI (Kadota et al. 2020). The library was sequenced on a HiSeq X sequencing platform (Illumina Inc., San Diego, CA, USA). The Hi-C reads were processed using Trim Galore! v0.6.8 (https://www.bioinformatics.babraham.ac.uk/projects/trim_galore/) with default settings and aligned to the HiFi contigs using Juicer ver. 1.6. Scaffolding of the primary unitigs was performed using 3D-DNA ver. 201008 (Dudchenko et al. 2017), applying the following parameters: ‘-m haploid -i 5000 -r 0--editor-repeat-coverage 20--editor-coarse-resolution 2500000--editor-coarse-region 12500000--editor-coarse-stringency 1--polisher-coarse-resolution 2500000--polisher-coarse-region 15000000--polisher-coarse-stringency 1--splitter-coarse-resolution 2500000--splitter-coarse-region 15000000--splitter-coarse-stringency 1’, as described by Tanaka et al. (2023). The remaining structural errors were manually corrected by inspecting the Hi-C contact map using Juicebox Assembly Tools (**Figure S1**). During this process, we also evaluated false contact signals along the diagonal, which may result from sequence similarity between paired scaffolds of homologous chromosomes. Scaffolding yielded chromosome-scale assemblies corresponding to the 36 chromosomes. For each homologous pair, the longer scaffold was designated as the primary chromosome and the shorter as the alternative chromosome. These were then organized into the “primary” and “alternative” assemblies, respectively. Assembly continuity and completeness, along with the predicted protein-coding gene set, were assessed using the gVolante webserver ver. 2.0.0 (Nishimura et al. 2017), which implements the Benchmarking Universal Single-Copy Orthologs (BUSCO) pipeline ver. 5 (Manni et al. 2021), using the ‘Metazoa’ or ‘Mollusca’ ortholog sets.

### Repeat identification and protein-coding gene prediction

The primary and alternative assemblies were subjected to *de novo* repeat element identification using RepeatModeler ver. 2.0.4 (Flynn et al. 2020). Identified repeat elements were used to mask both assemblies using RepeatMasker ver. 4.1.6 (Smit et al. 2015). These analyses were conducted using the Dfam TE Tools Container ver. 1.88 (https://github.com/Dfam-consortium/TETools). Protein-coding gene prediction was performed using Braker ver. 2.1.5 (Brůna et al. 2021), an automated pipeline for *ab initio* gene prediction that integrates RNA sequencing and protein homology evidence. RNA-seq data for *H. gigantea* were downloaded from the NCBI Sequence Read Archive (SRA, Leinonen et al. 2011) using the following accessions: SRR1910554, SRR1943392, SRR1944101, SRR1944137, SRR1985199, SRR1985200, SRR14327194, SRR14327195, SRR14327196, SRR14327197, SRR14327202, SRR14327203, SRR14327204, SRR14327205, SRR14327206, SRR14327207, SRR14327208, SRR14327209. All reads were aligned to the *H. gigantea* genome assembly as paired-end reads using HISAT2 ver. 2.1.0 (Kim et al. 2019). Gene prediction was carried out on the soft-masked primary and alternative assemblies by integrating the RNA-seq BAM files and protein sequences from two abalone genomes (*H. disucs hannai*: Nam et al. 2017; *H. rufescens*: Masonbrink et al. 2019), along with metazoan protein sequences from OrthoDB ver.10 (metazoa_proteins.fasta: Kuznetsov et al. 2023). Candidate coding sequences predicted by Braker2 were further analyzed using TransDecoder (http://transdecoder.sourceforge.net/) to identify open reading frames. Predicted coding sequences were functionally annotated using InterProScan ver. 5.14-53.0 (Jones et al. 2014). Only coding genes with identified functional domains were retained and used as the final gene sets for both assemblies to reduce false positives in gene prediction. The completeness of these gene sets was assessed using gVolante with the BUSCO ver. 5 pipeline and the ‘Metazoa’ ortholog set. Analyses were performed using default settings. For each predicted gene, the longest protein isoform was obtained.

### Comparison between primary and alternative assemblies

Pairwise alignment of the primary and alternative assemblies was performed using NUCmer ver. 4.0.0 (Marçais et al. 2018). The resulting file was filtered using the Delta-filter tool (−1 -i 70 −l 1000) and converted to a coordinate file using Show-coords tool. Dot plots were generated from the coordinate file using DotPlot (https://github.com/tpoorten/dotPlotPlot) to visualize collinearity between assemblies. Syntenic regions and structural rearrangements between the primary and alternative assemblies were identified using SyRI ver. 1.5.4 (Goel et al. 2019), based on pairwise alignments generated by minimap2 ver. 2.24 (-ax asm5). The results from SyRI were visualized using PlotSr ver. 0.5.3 (Goel and Schneeberger 2022). To identify synteny blocks between the two assemblies, we used MCScanX (Wang et al. 2012). MCScanX was run with default parameters, requiring a minimum of five genes to define a syntenic block. Protein sequences from the two assemblies were compared using BLASTP (*E* value <1 × 10^−10^), and the resulting alignments were used as input for MCscanX.

### Genomic features among chromosomes in primary and alternative assembly

We examined the distribution of eight genomic features across scaffolds in the primary and alternative assemblies: counts of BUSCO genes, protein-coding genes, repeat elements, tandem duplication genes, segmental duplication genes, and structural variations (deletions, insertions, and inversions). Tandem and segmental duplication genes were identified using the duplicate_gene_classifier tool implemented in MCScanX. Structural variations were detected using pbsv ver. 2.6.2 (https://github.com/PacificBiosciences/pbsv), with HiFi reads aligned to the primary assembly using pbmm2. The pbsv Discovery module was used to detect signatures of structural variations, including deletions, insertions, and inversions, and to call SVs. The number of each genomic feature within the 5 Mb sliding-window was quantified using the intersected function in bedtools (Quinlan and Hall 2010). Feature counts across chromosomes were compared using analysis of variance (ANOVA), followed by Tukey’s post hoc honest significant difference test.

### Comparison between the primary assembly and other abalone assemblies

We compared the *H. gigantea* primary assembly with five previously published abalone genome assemblies: *H. cracherodii* (Orland et al. 2022), *H. rufescens* (Griffiths et al. 2022), *H. rubra* (Gan et al. 2019), *H. midae* (Tshilate et al. 2023), and *H. asinina* (Barkan et al. 2024). Pairwise alignments were performed using NUCmer, as described above. The resulting coordinate files were used to calculate the alignment percentages of each abalone genome against *H. gigantea* chromosomes. For *H. cracherodii* and *H. rufescens*, both primary and alternative scaffolds corresponding to homologous chromosomes are available (Orland et al. 2022; Griffiths et al. 2022), allowing for comparison of their two haplotype assemblies as described for *H. gigantea*.

### Domain enrichment analysis

Pfam domains enriched in the three non-syntenic homologous chromosomes (*see the Result section*) were identified using InterProScan, and statistical enrichment was assessed using a hypergeometric test, corrected for multiple testing using the Benjamini–Hochberg method in R. To identify gene names associated with the most enriched Pfam domains in each non-syntenic homologous chromosome, we performed BLASTP searches (*E* value <1 × 10^−8^) against *H. rufescens* protein sequences obtained from the Ensembl Metazoa database.

### Alignment depth analysis

Whole-genome sequencing data from eight individuals each of *H. discus* and *H. madaka*, and six individuals of *H. gigantea*, were used for alignment depth analysis. These data were generated using the Illumina HiSeq X Ten platform with 150-bp paired-end reads, as described in our previous study (Hirase et al. 2021). Raw reads were filtered using Trimmomatic ver. 3.8 (Bolger et al. 2014) to remove adapters, Illumina-specific sequences, and low-quality regions using the parameters in Hirase et al. (2021). Filtered paired-end reads were aligned to the *H. gigantea* primary assembly using the BWA-MEM algorithm (Li and Durbin 2009; Li 2013) with default settings. Local realignment around indels was performed using GATK ver. 3.7 (McKenna et al. 2010), also with default settings.

### Proportion of duplicated genes of fully sequenced invertebrates

To evaluate the abundance of duplicated genes in abalone species, we estimated the proportion of duplicated genes (*P*D) in 46 invertebrate species, including four abalone species (**Table S1**). Amino acid sequences for the invertebrate species other than abalone species were analyzed in previous studies (Makino and Kawata, 2019; Sato et al. 2023). Amino acid sequences for two abalone species (*H. rubra* and *H. rufescens*) were downloaded from the Ensembl Metazoa database. In addition, amino acid sequences for one abalone species (*H. asinina*) were obtained from Barkan et al. (2024). Duplicated genes were identified by performing an all-to-all BLASTP search of protein sequences within each species, following the method described in Makino and Kawata, 2019. Genes were considered duplicated if they had a homolog within the same species with an E-value < 10^-4^ and a query coverage > 30%. Genes annotated as transposons were removed using annotations from the PANTHER database ver. 11 (Mi et al. 2013). The number of protein-coding genes was then counted after excluding transposons, and PD was calculated as the proportion of duplicated genes among total protein-coding genes (excluding transposons; **Table S1**). Although PD has not been shown to correlate with BUSCO scores, a commonly used measure of genome completeness (Makino and Kawata, 2019; **Table S1**), we excluded published genomes with low BUSCO completeness (BUSCO < 80%) from our analysis to ensure accuracy.

## Results

### Genome sequencing, assembly, and gene prediction

HiFi long-read sequencing of the *H*. *gigantea* genome produced a total of 55.5 gigabases (Gb) of data, representing approximately 30.8x coverage of the genome, assuming a genome size of 1.8 Gb (Nam et al. 2017), with an average read length of 16.2 kilobases (Kb) (**Table S2**). These reads were assembled into 7,407 unitigs, yielding an N50 length of 2.19 megabases (Mb) and a maximum sequence length of 10.39 Mb (**Table S3**). The total unitig length was 2.8 Gb, compared to an estimated total genome length of 3.6 Gb, suggesting approximately 0.8 Gb of unassembled regions. Despite this, a high proportion (95.3%) of duplicated Metazoa BUSCO genes were detected, indicating that the diploid genome assembly is highly complete (**Table S3**). A Hi-C library was independently prepared and sequenced, generating 66.8 Gb of paired-end read data (**Table S2**). Hi-C scaffolding produced a chromosome-scale assembly comprising 4,534 scaffolds with an N50 of 71.2 Mb and a maximum scaffold length of 96.6 Mb (**Table S4**).

The chromosome-scale genome assembly included 36 scaffolds larger than 5 Mb, consistent with the diploid chromosome number of *H. gigantea* (2*n* = 36) (Miyake et al. 1997). This assembly was divided into two haplotype assemblies–primary and alternative–based on the distribution of Metazoa BUSCO genes. Chromosomal scaffold length differences between haplotypes ranged from 3.4 Mb (Chr6) to 0.08 Mb (Chr11), with an average difference of 1.3 Mb (**Table S5**). The primary assembly contained 96.5% complete Metazoa and 84.8% complete Mollusca BUSCO genes, whereas the alternative assembly contained 96.2% and 84.9%, respectively (**Table S4**). These values were comparable to those reported for other chromosome-level abalone assemblies (**Table S6**). Based on the gene order of Mollusca BUSCO genes, conserved synteny was observed between homologous haplotype pairs (**Figure 1)**. Collectively, these findings suggest that the 18 pairs of chromosomal scaffolds represent a high-quality, nearly complete haplotype-phased reconstruction of the *H. gigantea* genome. Gene prediction using Braker2 identified 62,479 and 61,384 total genes in the primary and alternative assemblies, respectively, of which 52,015 and 51,897 were protein-coding genes (**Table S7**). After functional domain filtering using InterProScan, 30,857 protein-coding genes were retained in the primary assembly and 34,896 in the alternative assembly. These final gene sets included more than 92% complete Metazoa and over 82% complete Mollusca BUSCO genes (**Table S7**).

**Figure 1.**
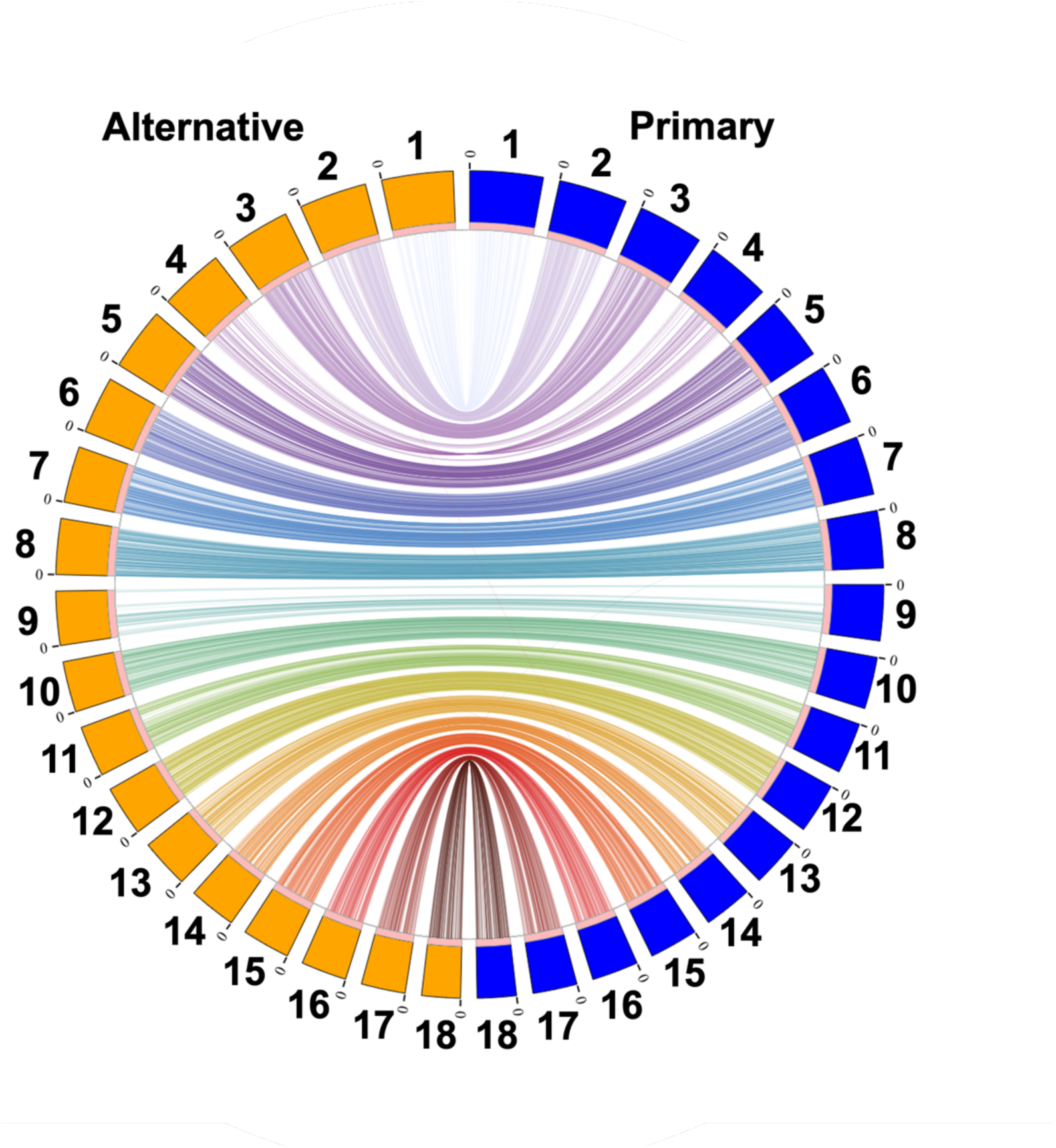
Synteny plot between primary (blue) and alternative (orange) assemblies based on Mollusca Benchmarking Universal Single-Copy Orthologs (BUSCO) genes.

### Three non-syntenic homologous chromosomes

We compared the primary and alternative assemblies based on nucleotide sequence alignments. Collinearity of sequences was generally conserved across most chromosomes, although several large inversions were detected (**Figure 2**). In particular, the Hi-C contact map suggested that a large inversion on Chr4 was unlikely to be an assembly artifact (**Figure S1**). Notably, the percentage of aligned genomic regions between pairs of homologous chromosomes was less than 85% for Chr1, Chr4, and Chr9 (**Figure 3**), and these chromosomes also showed extensive duplicated and rearranged regions (**Figure 2**). Henceforth, we refer to these three chromosomes as “non-syntenic homologous chromosomes”. Even when compared with genome assemblies from the closely related North American abalone species *H. cracherodii* and *H. rufescens*,alignments for these chromosomes could not be achieved well (**Figure 3, S2)**. While we considered the possibility that misassemblies in the primary or alternative assemblies may have contributed to the observed non-syntenic patterns, similar patterns were also detected between the haplotype-resolved scaffolds *H. cracherodii* and *H. rufescens* (**Figure S3, S4**). This observation suggests that the non-syntenic features are not specific to misassembly in this study and may reflect genuine evolutionary genomic changes.

**Figure 2.**
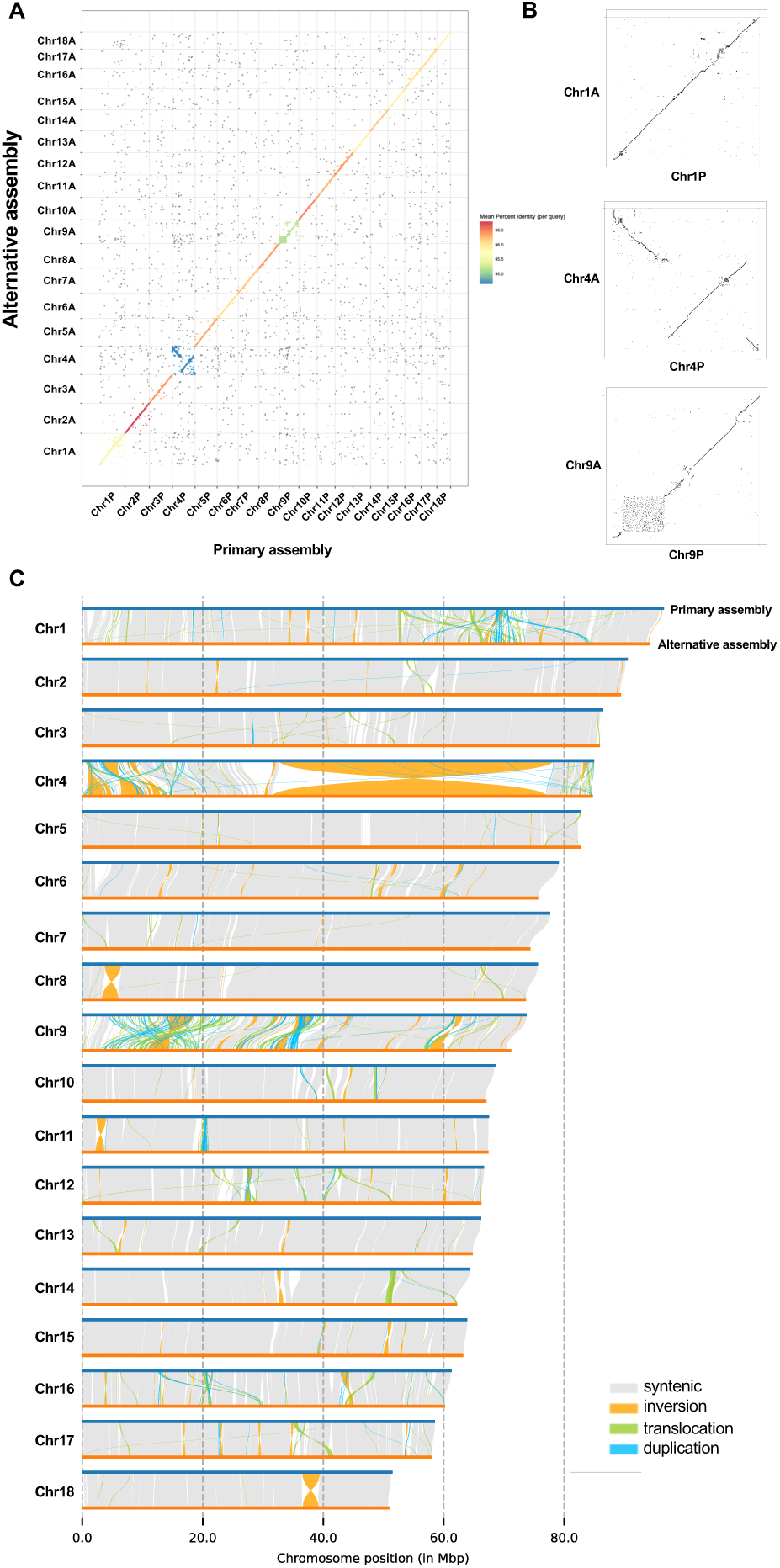
Genome-wide syntenic relationships between the primary (Chr1P–Chr18P) and alternative (Chr1A–Chr18A) assemblies. (A) Dot plot of alignment between the primary and alternative assemblies across all chromosomes, with mean percent identity per query shown by color. (B) Dot plot of alignment for the three non-syntenic chromosomes. (C) Synteny and rearrangement (SyRI) plot comparing the primary and alternative assemblies.

**Figure 3.**
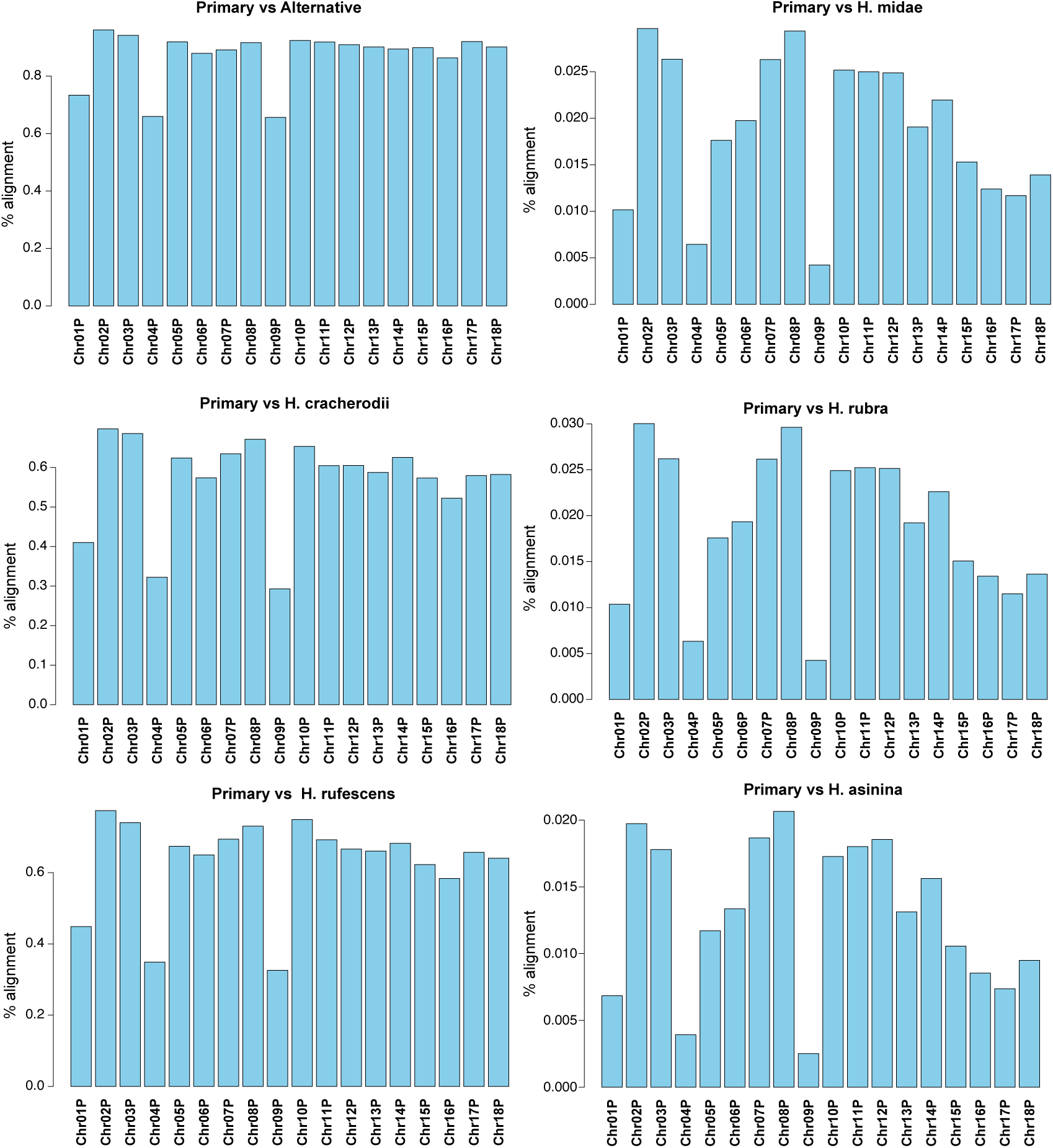
Percentage of genomic regions in the alternative assembly and other abalone assemblies aligned to the primary assembly (Chr1P–Chr18P). *Haliotis cracherodii* and *H. rufescens* are closely related to *H. gigantea*, whereas *H. rubra, H. midae, and H. asinina* are distantly related.

### The non-syntenic patterns generated by multiple gene duplication events

We examined the distribution of BUSCO genes, protein-coding genes, tandem duplication genes, segmental duplication genes, repeats, deletions, insertions, and inversions across chromosomes (**Figure 4; FigureS5-16; Table S8-21**). Three distinct genomic patterns were observed in the non-syntenic homologous chromosomes. First, the number of BUSCO genes was significantly lower in these chromosomes relative to the total number of genes (*P* < 0.05; **Figure 4; Table S5, S6, S11, S12**). Second, an uneven distribution of BUSCO genes, segmental duplication genes, and repeat elements were found in Chr9 (**Figure S5, S7, S8**). Third, Chr4 showed a high accumulation of deletions and insertions (**Figure S10, S16; Table S19, S20**). The region with many insertions and deletions in Chr4 overlapped with the location of the large inversion, suggesting a potential association between these events (**Figure 2, S10**).

**Figure 4.**
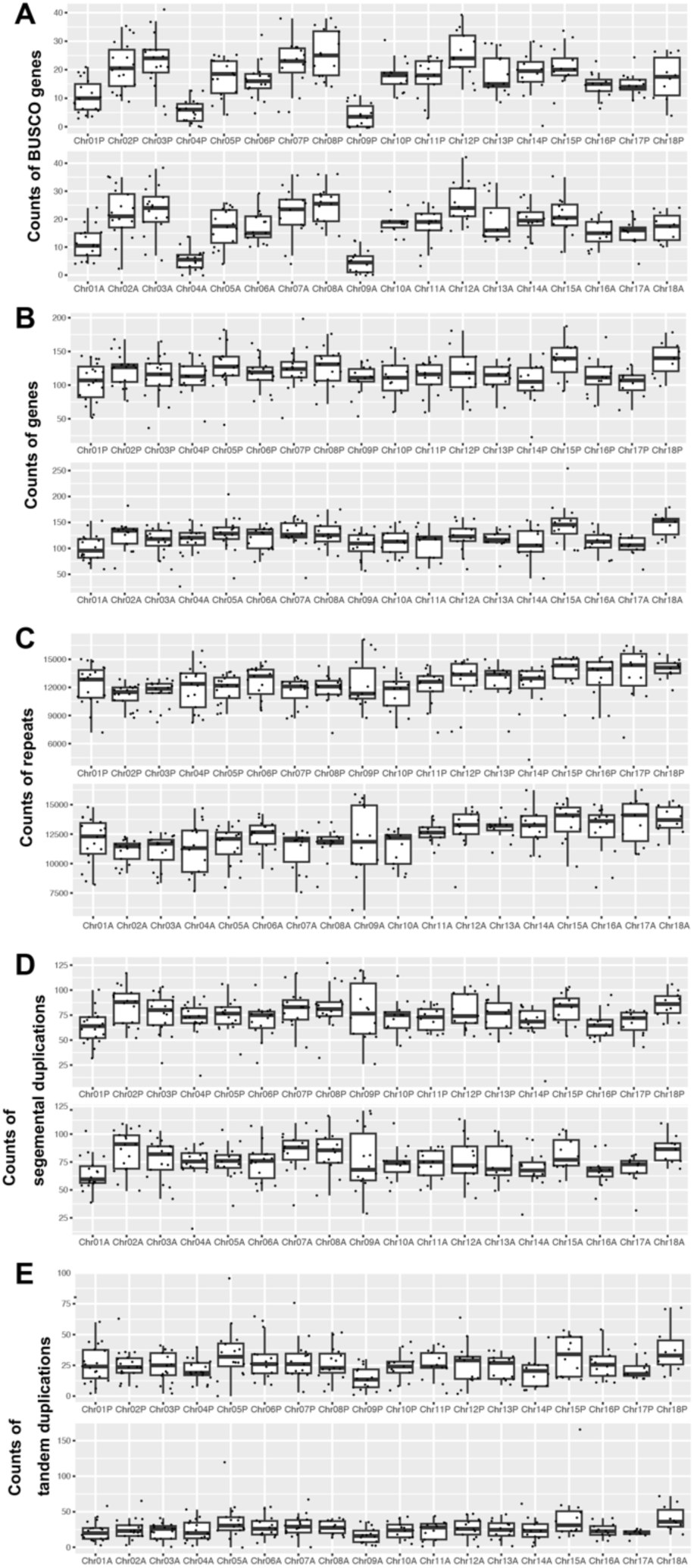
Boxplots showing the distribution of five genomic features across chromosomes in the primary and alternative assemblies, calculated using 5-Mb sliding windows: (A) counts of Metazoan benchmarking universal single-copy orthologs (BUSCO) genes, (B) coding genes, (C) repeats, (D) segmental duplication genes, and (E) tandem duplication genes.

We used MCscanX to assess interchromosomal and intrachromosomal syntenic patterns by identifying syntenic gene blocks comprising at least five genes (**Figure 5; Figure S17-19)**. Numerous syntenic blocks were detected within each of the three non-syntenic homologous chromosomes, indicating that frequent segmental duplications occurred in these regions (**Figure 5**). The dot plot of primary and alternative assemblies revealed asymmetry in the arrangement of syntenic blocks, particularly in the non-syntenic homologous chromosomes (**Figure 5A**), further supporting differential duplication events between the haplotypes. Many syntenic gene blocks were shared between Chr1 and Chr9, suggesting segmental duplication exchange or parallel amplification between these non-syntenic homologous chromosomes. These syntenic block patterns were also detected in *H. rufescens* scaffolds corresponding to the same three homologous chromosomes (**Figure 5; Figure S20**), implying that these structural characteristics may be conserved across multiple abalone species.

**Figure 5.**
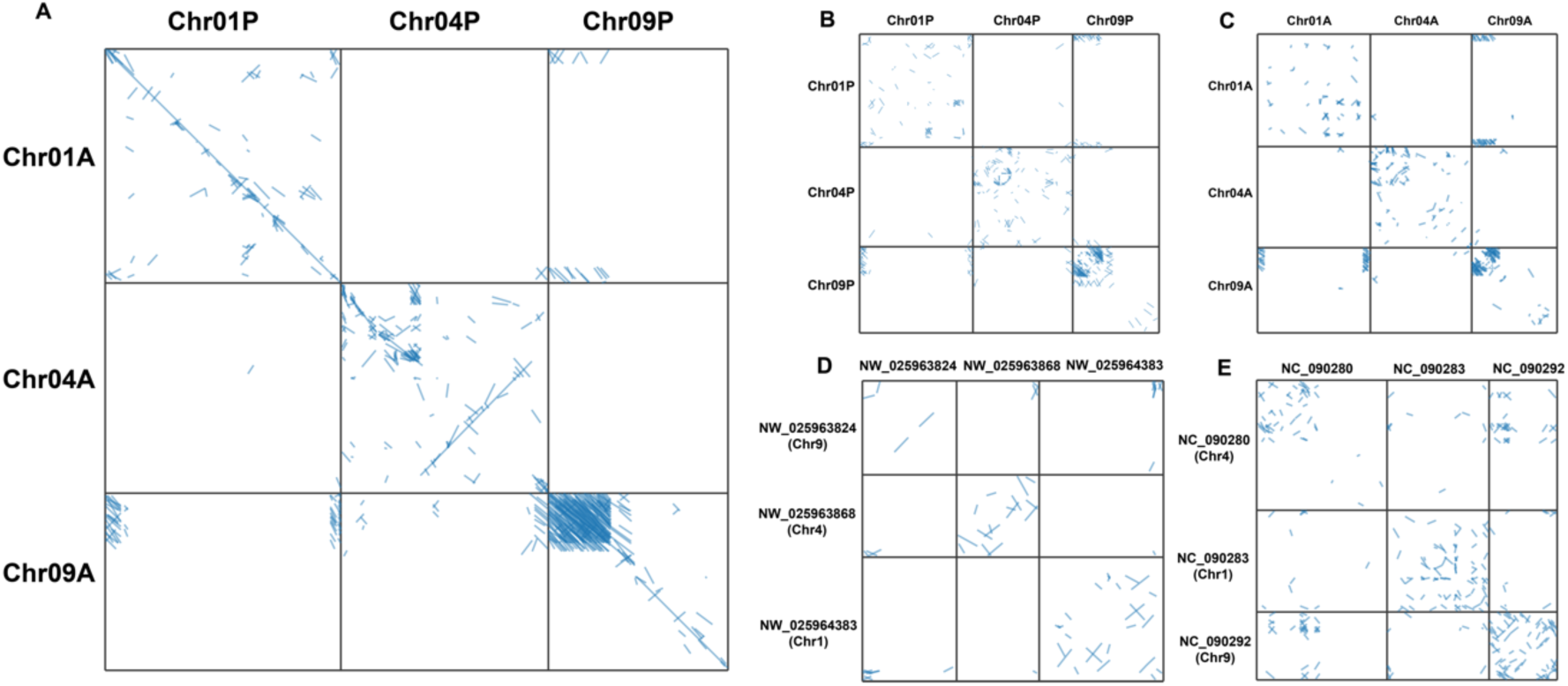
Dot plots of interchomosomal and intrachomosomal syntenic gene blocks for the three non-syntenic chromosomes in the primary and alternative assemblies (A-C) and in scaffolds of *Haliotis rufescens* (D) and *H. asinina* (E).

### Ancestral origin of non-syntenic homologous chromosomes in abalones

Globally, abalones can be divided into two major evolutionary lineages: one includes species from the Japanese Archipelago and the North American coast, while the other includes species from other regions, such as *H. rubra* and *H. asinina* from Australia and *H. midae* from South Africa (Streit et al. 2006). Chromosome numbers differ between these two lineages (2*n* = 36 vs. 2*n* = 32) (Arai and Okumura 2013), suggesting that significant genomic changes have occurred. In the latter lineage, *H. asinina* is the only species for which chromosome-level genome assemblies are currently available. Based on our analysis of the *H. gigantea* genome, we found that the difference in chromosome number could be explained by chromosome fusion or fission events involving Chr4,Chr11, Chr15, and Chr18 (**Figure S2).**

Regions of the three non-syntenic homologous chromosomes in *H. gigantea* showed less homology to any regions of the *H. rubra*, *H. asinina*, or *H. midae* genomes than those of other chromosomes (**Figure 3**; **Figure S2**). Notably, patterns of intrachromosomal segmental duplication patterns were detected in three *H. asinine* scaffolds that showed orthologous relationships with the three non-syntenic homologous chromosomes of *H. gigantea*: NC_090283 with Chr1, the first half of NC090280 with Chr4, and NC_090292 with Chr9 (**Figure 5E; Figure S21)**.

### Enrichment of immune-related domains in non-syntenic homologous chromosomes

We observed functional domain enrichment in the three non-syntenic homologous chromosomes: the major facilitator superfamily (MFS) general substrate transporter domain was enriched in Chr1 (**Figure 6A, 6B, Table S22, S23**), the immunoglobulin domain in Chr4 (**Figure 6C, 6D, Table S24, 25**), and the ankyrin repeat domain in Chr9 (**Table S26, 27**). These domains were also enriched in the corresponding *H. asinine* chromosomes, orthologous to the non-syntenic homologous chromosomes in *H. gigantea* (**Figure 6E, 6F, Table S28, S29, S30**).

**Figure 6.**
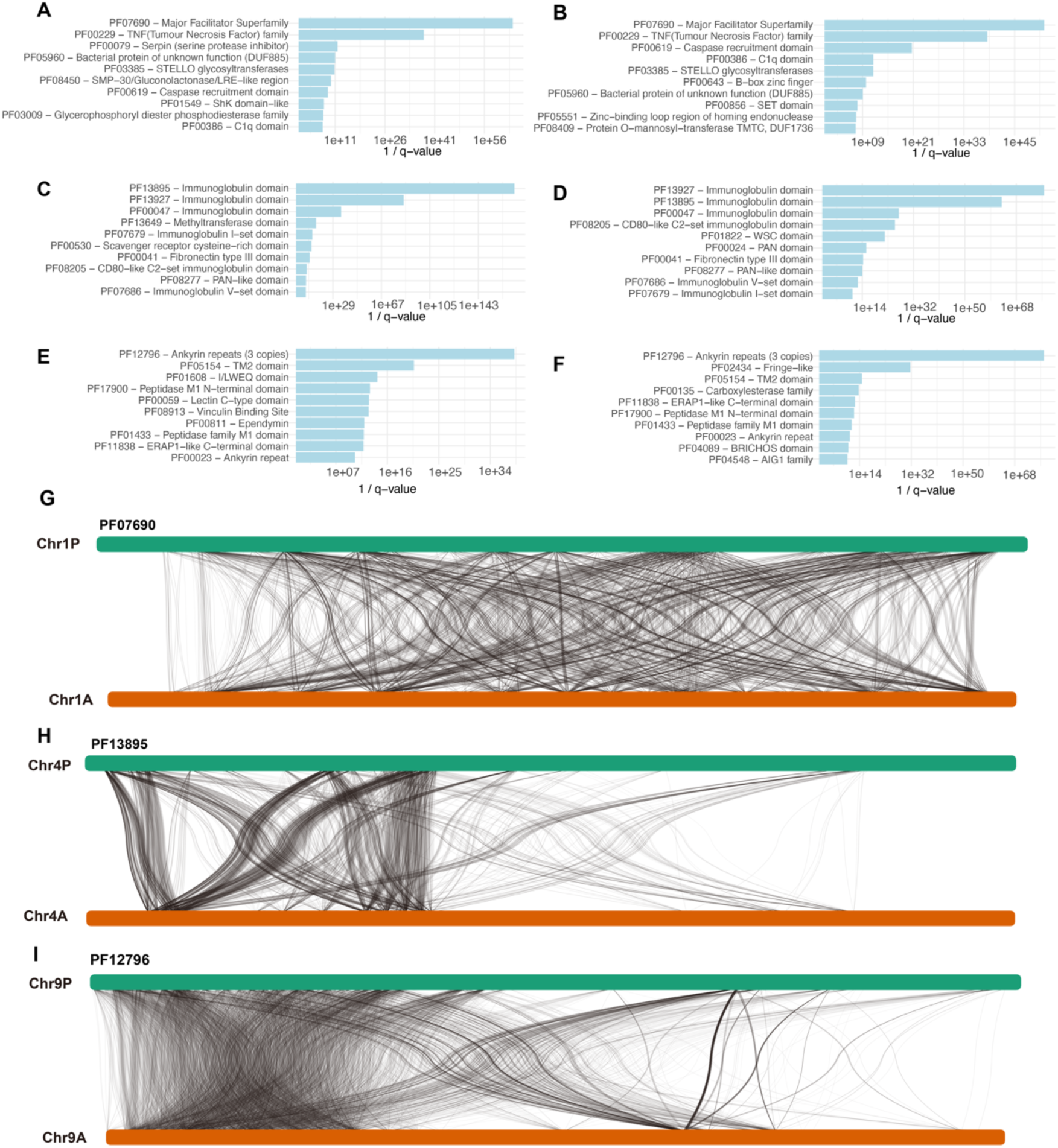
Top 10 enriched domains in Chr1P (A), Chr1A (B), Chr4P (C), Chr4A (D), Chr9P (E), and Chr9A (F). Synteny plots showing duplication relationships of genes with the top enriched domains in Chr1 (G), Chr4 (H), and Chr9 (I).

On Chr1, genes with MFS domains include members of the solute carrier (SLC) family and monocarboxylate transporters. Chr4 contains immunoglobulin domain-bearing genes, such as those encoding neural cell adhesion molecules. Chr9 is enriched with ankyrin repeat-containing genes, including those encoding ankyrin proteins and serine/threonine protein phosphatases. Analysis of the primary assembly confirmed that genes with these functional domains were particularly abundant on their respective chromosomes (**Figure S22**). Additionally, localized increases in genes with MFS and ankyrin repeat domains were also detected on other chromosomes, although their enrichment was most prominent in Chr1 and Chr9, respectively.

### Accumulation of duplicated genes in abalones

The proportion of duplicated genes (*P*D) has been reported to be higher in species with smaller propagule sizes, defined as the size of eggs or juveniles dispersed from their parents (Makino and Kawata 2019). Abalones are known to have relatively small propagule sizes (Sherman et al. 2020; Phan et al. 2022, Table S1). In addition, our findings demonstrate the presence of non-syntenic chromosomes shared across abalone species. These combined observations may help explain the notably high number of duplicated genes observed in abalones compared to other invertebrates. As expected, we found high *P*D values in abalone species among the 46 invertebrates examined (**Figure 7**). Moreover, *P*D values were negatively correlated with propagule size across invertebrates (Pearson correlation coefficient, *r* = −0.547, *P* < 0.001), consistent with previous studies (Makino and Kawata 2019). Given their relatively small propagule size, the high *P*D values observed in abalones are not surprising. However, since *P*D values in abalones were elevated even relative to invertebrates with similarly small propagule sizes, the presence of non-syntenic chromosomes likely contributed to the increased gene duplication observed in abalone genomes.

**Figure 7.**
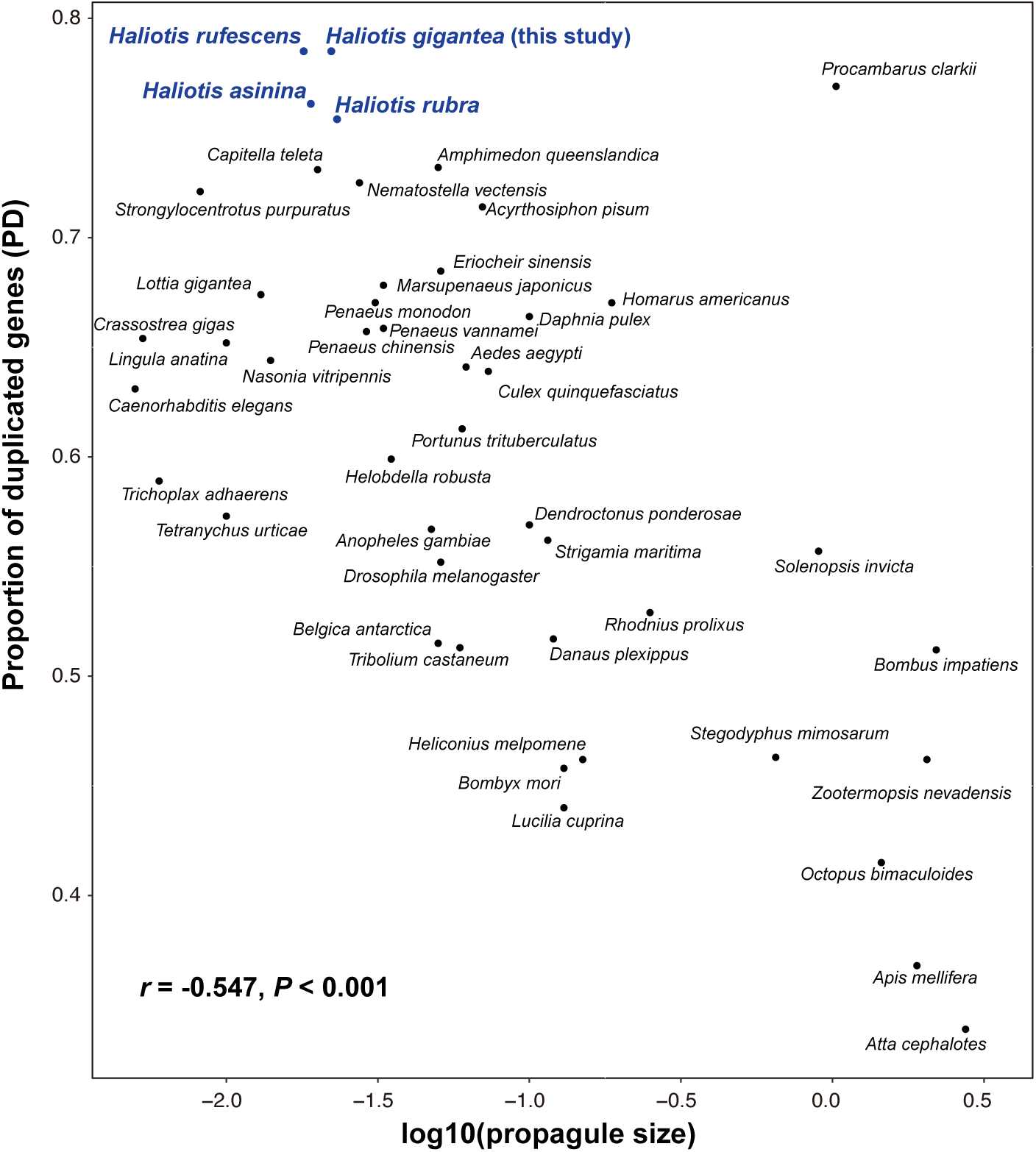
Relationship between the proportion of duplicated genes (*P*D) and propagule size of fully sequenced 46 invertebrate species including four *Haliotis* species.

### Variation in sequence depth between individuals and species in homologous chromosomes

Short-read sequencing data from regional populations of *H. gigantea* and two closely related species, *H. disucs* and *H. madaka* were mapped to the *H. gigantea* primary genome assembly. Read coverage, defined as the proportion of regions where short reads were mapped, was consistently lower in non-syntenic homologous chromosomes compared to other chromosomes (**Figure S23A**). This reduction in coverage was attributed to a higher proportion of multi-mapped reads in the non-syntenic homologous chromosomes (**Figure S23B**). Bartlett’s test revealed significant heterogeneity in read depth variance among chromosomes (*P* < 0.05). Furthermore, both the average read depth and inter-individual variance were higher in non-syntenic homologous chromosomes than in syntenic chromosomes across all three species when multi-mapped reads were included (**Figure S24-S26**). These results suggest the presence of copy number variations in the three homologous chromosomes and support their genomic instability not only in *H. gigantea* but also in closely related western Pacific abalone species.

## Discussion

We constructed a haplotype-phased genome assembly of *H. gigantea* and identified three of the 18 homologous chromosomes that exhibited non-syntenic patterns. These chromosomes did not align well with those of other abalone species, and they contained many segmental duplications. Similar non-syntenic regions, resulting from gene duplication, have been reported in plant species (Jiao and Schneeberger 2020; Sun et al. 2022) and marine invertebrates (Takeuchi et al. 2022). However, the non-syntenic regions identified in *H. gigantea* were larger than those observed in the previous studies and extended across greater portions of each chromosome. These findings suggest that the presence of such chromosomes contributes to the unusually high number of duplicated genes in abalone genomes. Notably, segmental duplications occurred more frequently between non-syntenic homologous chromosomes than between other chromosomal regions. This pattern may indicate the existence of a mechanism that facilitates gene duplication more among these specific chromosomes. The number of BUSCO genes was low in the non-syntenic chromosomes. Since BUSCO genes typically occur as single copies per species (Manni et al. 2021), this observation suggests that the non-syntenic homologous chromosomes contain relatively few single-copy genes and a high number of duplicated genes. Universal single-copy genes are generally under strong purifying selection, as imbalances in their expression levels caused by duplication can have deleterious effects (Makino et al. 2013). Therefore, the scarcity of BUSCO genes in these chromosomes may be related to the evolutionary processes underlying their origin. Although non-syntenic regions are often associated with sex chromosomes in animals, none of the three non-syntenic homologous chromosomes identified in this study were associated with phenotypic sex (Kina et al. 2024). Further studies are needed to clarify the evolutionary significance and biological roles of these chromosomes.

*Haliotis gigantea* and its closely related species in the Japanese Archipelago are thought to have diverged from amphi-Pacific abalones that radiated along the California coast during the glacial cycles (Geiger and Groves 1999). This hypothesis is supported by previous phylogenetic studies, which suggest close genetic relationships between western Pacific and North American abalone species (Lee and Vacquier 1995; Streit et al. 2006; Hirase et al. 2021). The scaffolds corresponding to non-syntenic homologous chromosomes (Chr1, Chr4, and Chr9) in North American species did not show one-to-one alignment with those of *H. gigantea,* indicating that the non-syntenic chromosomal structures may have been established before the divergence of *H. gigantea* from its North American relatives. Furthermore, regions of the three non-syntenic homologous chromosomes in *H. gigantea* did not align with any genomic regions in more distantly related abalone species from South Africa and Australia. In *H. asinine,* segmental duplications were also detected in the chromosomes orthologous to these homologous chromosomes in *H. gigantea*. Although haplotype-phased genome assemblies for these distantly related species are currently unavailable, the presence of segmental duplications in orthologous chromosomes suggests that non-syntenic states may have originated in the early stages of abalone evolution, possibly during or soon after their emergence in the Cretaceous Period (Geiger and Owen 2012).

We found that specific functional domains were enriched in the three non-syntenic chromosomes of *H. gigantea* and the corresponding scaffolds of *H. asinina*, and that the types of enriched domains differed between them. By focusing on these domains in *H. gigantea* genome, we identified gene families that had undergone duplication in a chromosome-specific manner. In the Chr1 scaffold, many solute carriers (SLC) and monocarboxylate transporter (MCT) genes have an MFS general substrate transporter domain. Previous studies have suggested that SLC genes play critical roles in adaptation to salinity and temperature fluctuations in various marine taxa (Xu et al. 2013; Barrio et al. 2016; Zhou et al. 2018; Ertl et al. 2019; Zhao et al. 2021). In addition, several SLC family genes have been proposed as candidates involved in the speciation of *H. discus* and *H. madaka*, which exhibit different habitat depths (Hirase et al. 2021). The Chr4 scaffold included several neural cell adhesion molecule (NCAM) genes with immunoglobulin domains. These domains are also enriched in non-syntenic regions between homologous chromosomes in pearl oysters (Takeuchi et al., 2022). Takeuchi et al. (2022) hypothesized that duplicated immunoglobulin domains enhance immune gene repertoire and are important for innate immunity in oysters. Similarly, NCAM genes with immunoglobulin domains may contribute to disease resistance in *H. gigantea*. In the Chr9 scaffold, we identified duplicated ankyrin and serine/threonine protein phosphatase genes with ankyrin repeat domains. Transcriptome analysis of *H. midae* populations revealed an expansion of ankyrin repeat domains and suggested that this gene family is linked to host defense and may be important for survival in pathogen-rich environments (Picone et al. 2015). Furthermore, a comparison of gene expression between resilient and susceptible abalones indicated that ankyrin repeat-containing genes were significantly upregulated in resilient individuals, suggesting a potential role in enhancing preemptive defense mechanisms (Shiel et al. 2017).

This study has two limitations. The first is that there may be approximately 0.8 Gb of unassembled regions, so it cannot be said that the complete picture of *H. gigantea* genome has been fully elucidated. To effectively address this issue, it is likely necessary to obtain both longer and more abundant sequencing reads, as increased read length and coverage would significantly enhance the completeness and accuracy of the genome assembly. Second, although the haplotype-phased assembly revealed substantial structural variation and gene duplications, functional validation of the duplicated genes was not performed. The roles of these genes in environmental adaptation, immunity, or development remain to be confirmed through transcriptomic or experimental approaches. Additionally, haplotype-phased genome assemblies are still lacking for many abalone species, limiting the resolution of comparative genomic analyses across the entire genus. Future studies should aim to elucidate the functional consequences of gene duplications, particularly by examining their expression profiles under different environmental conditions or pathogen exposures.

## Acknowledgements

We express our gratitude to the Kikuchi Laboratory for their assistance. This study was supported by the Japan Society for the Promotion of Science (KAKENHI 23K05267 and 22H00377).

## Data accessibility

The genome assembly of *Haliotis gigantea* and the gene models associated to it, as well as the associated raw sequence reads, will be deposited in NCBI under the BioProject PRJDB20273 upon publication.

## Benefit-Sharing Statement

Benefits Generated: Although our research deals only with samples collected in Japan, the benefits from this research accrue from the sharing of our data and results in public databases, as described above.

## Author Contributions

SH conceived the study. SH collected samples. SH, MK, and SK designed the experiment and collected the data. SH, TM, and TT conducted the data analysis. SH wrote the manuscript with help from TM, TT, SK, and KK. All the authors approved the final version of the manuscript.

