## Supplementary figures for "Ancestral origin and structural characteristics of non-syntenic homologous chromosomes in abalones"

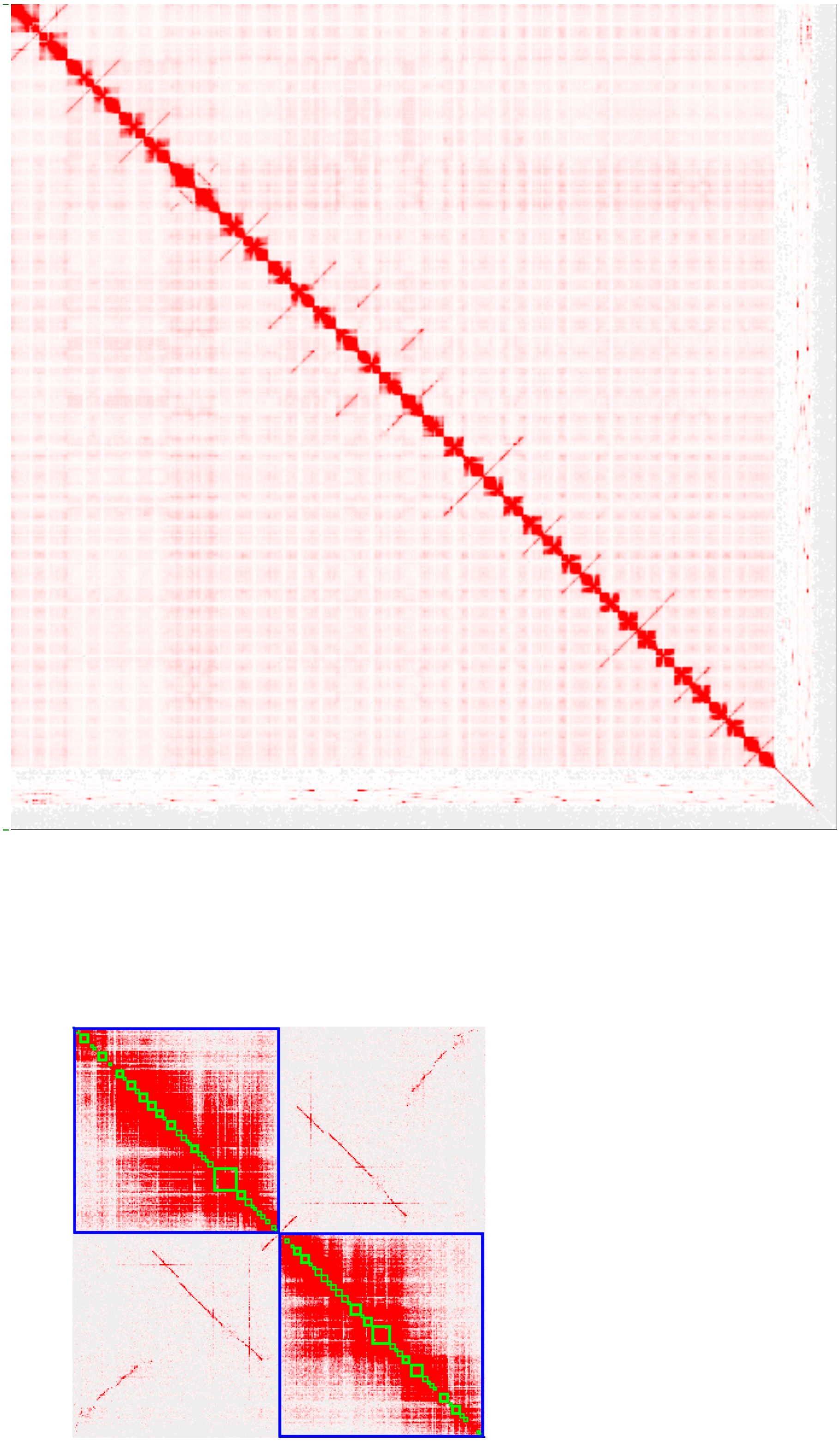

**Figure S1. A.** Genome contig contact matrix using Hi-C data, color bar indicates contact density from red (high) to white (low). **B.** Contact matrix for the primary and alternative Chr4 scafflolds.

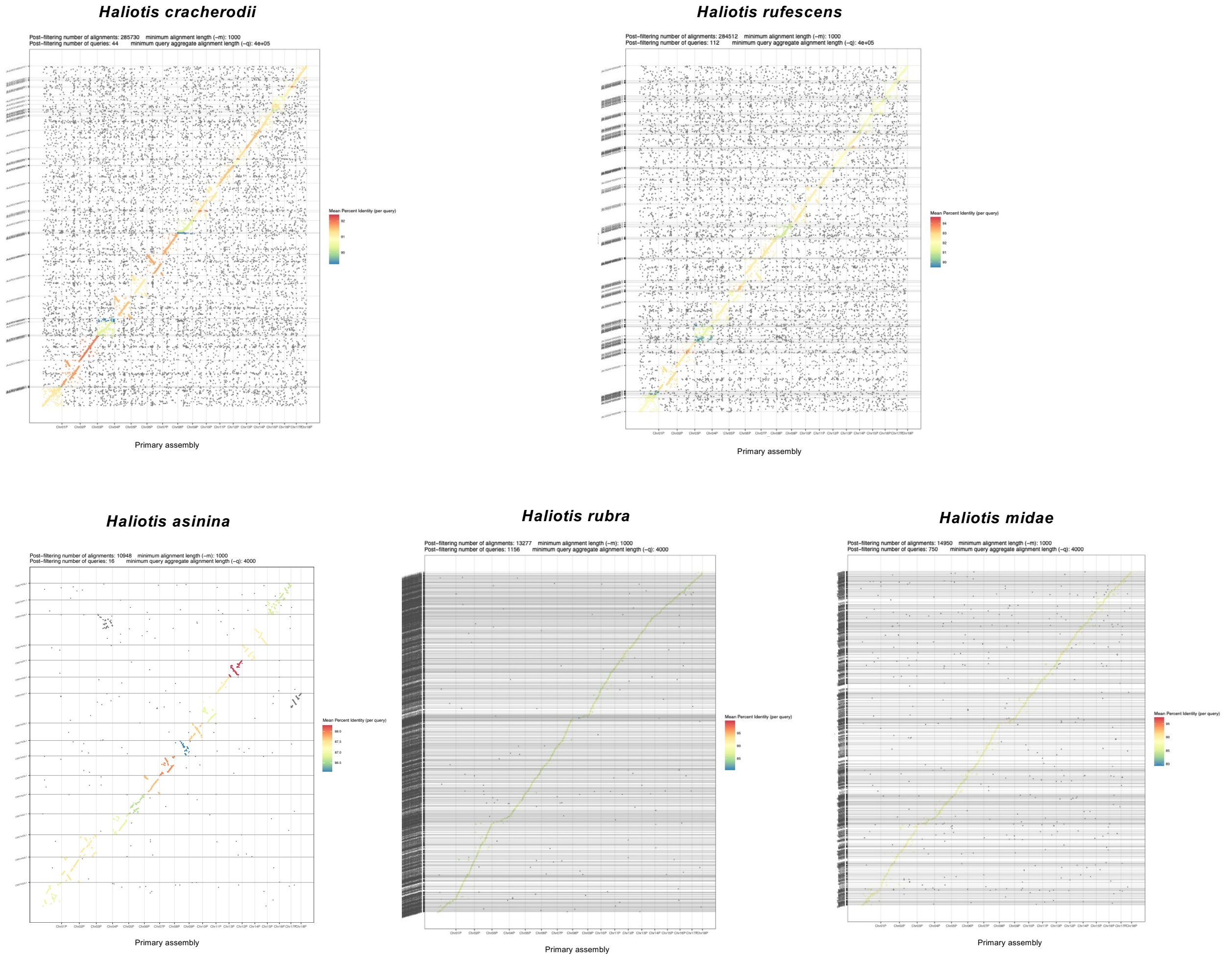


**Figure S2.** Dot plots of synteny relationship between the primary and other abalone genome assemblies. Mean percent identity per query was shown by color.

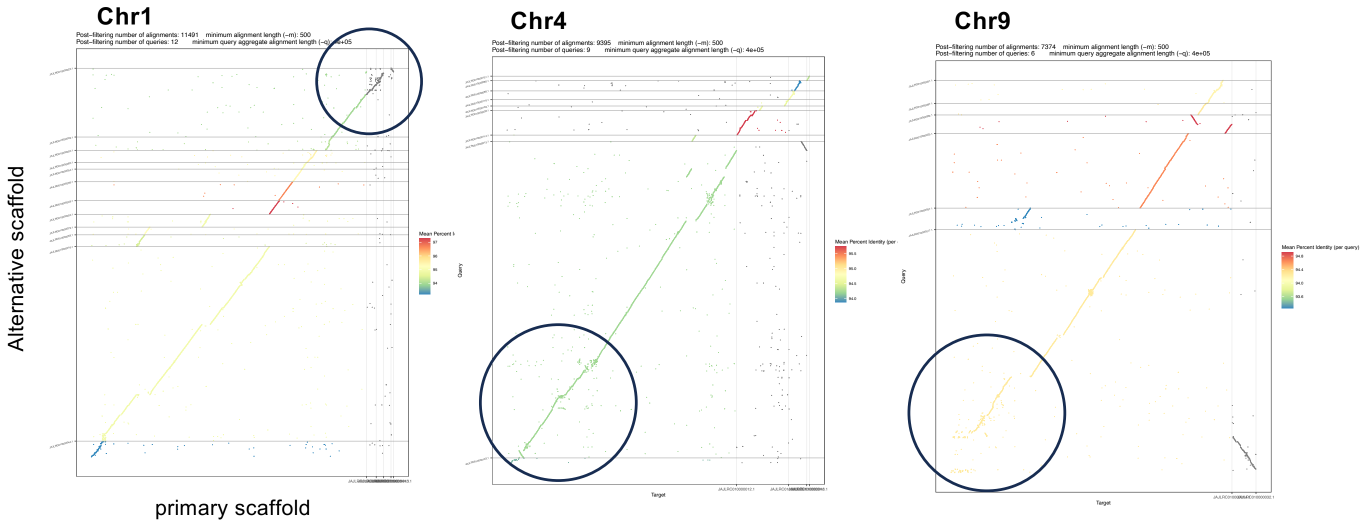


**Figure S3** Dot plots of synteny relationship between primary and alternative scaffolds of *Haliotis cracherodii* corresponding to Chr1, 4 and 9, respectively.

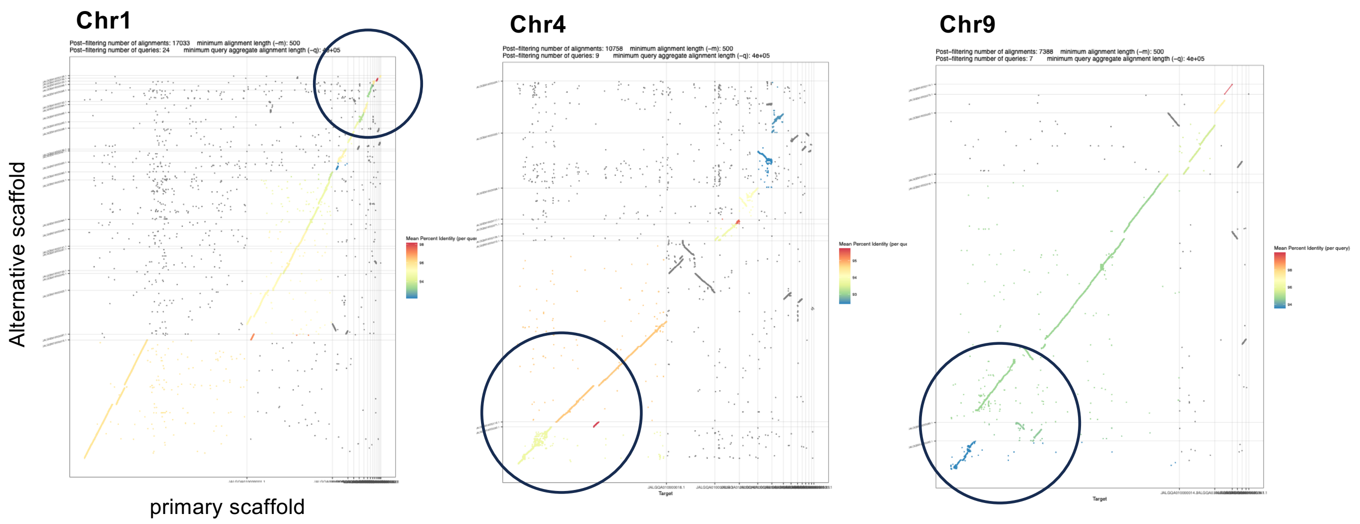


**Figure S4** Dot plots of synteny relationship between primary and alternative scaffolds of *Haliotis rufescens* corresponding to Chr1, 4, and 9, respectively.

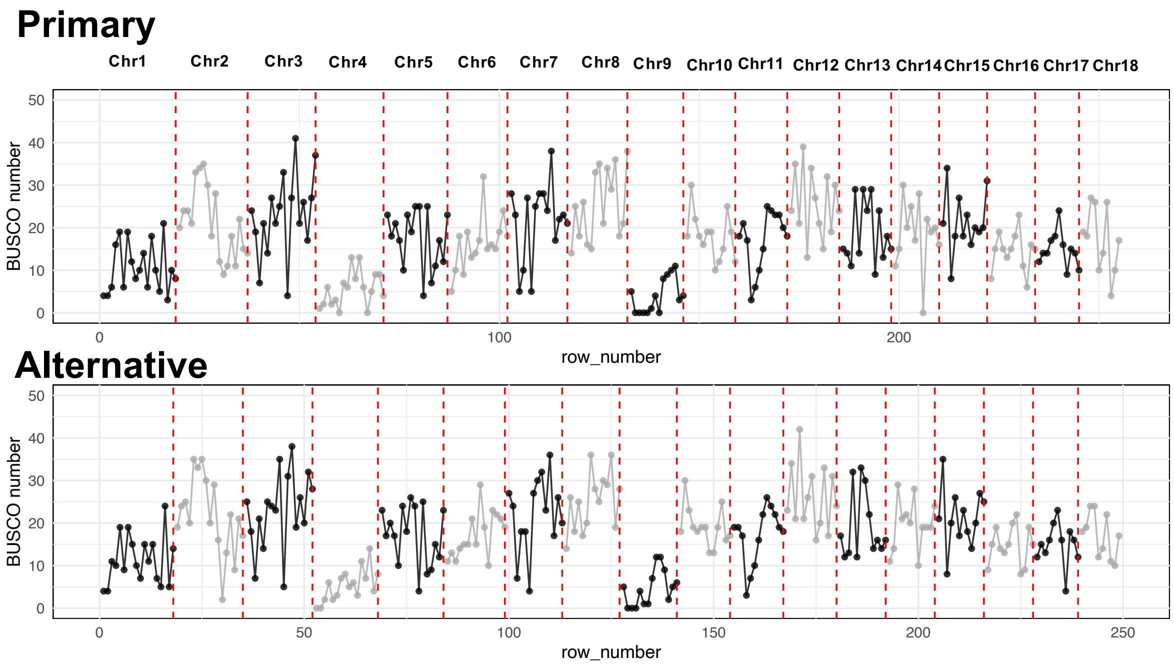


**Figure S5.** Counts of metazoa BUSCO genes in the primary and alternative assemblies using a 5 mega-base pair sliding window.

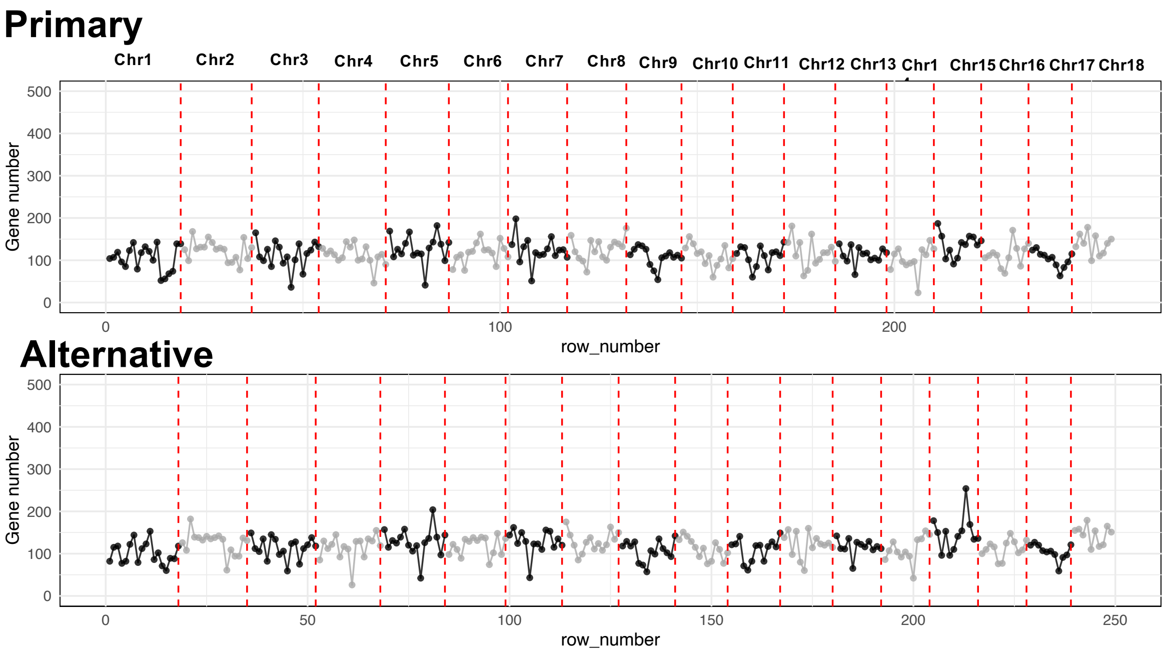


**Figure S6.** Counts of genes in the primary and alternative assemblies using a 5 mega-base pair sliding window.

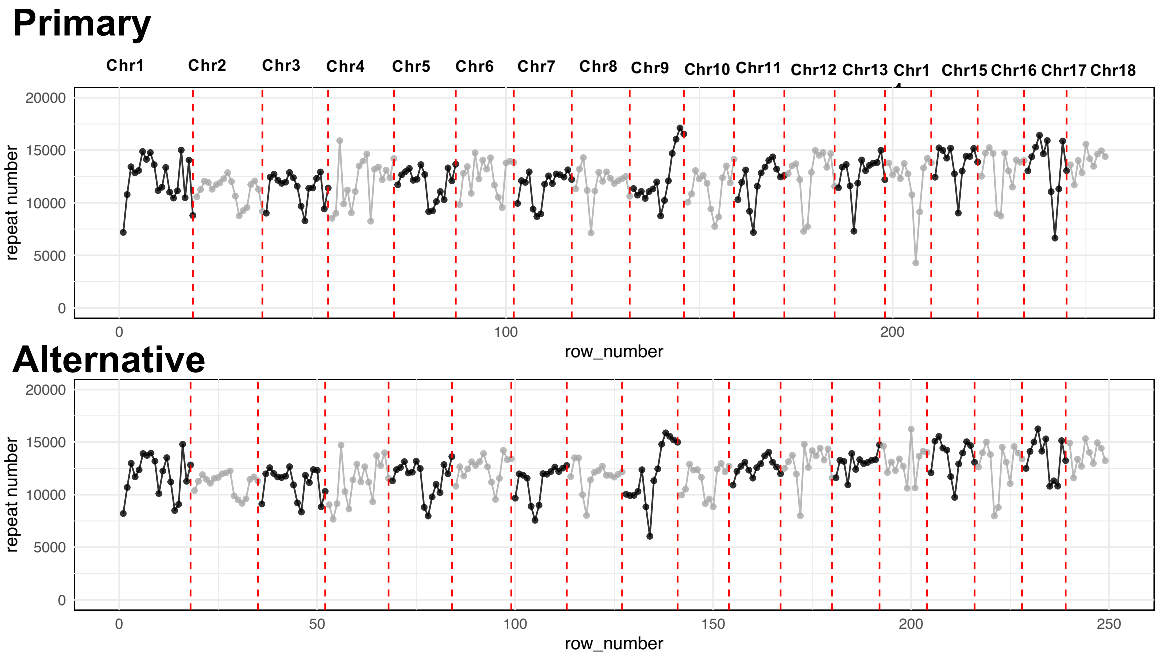


**Figure S7.** Counts of repeats in the primary and alternative assemblies using a 5 mega-base pair sliding window.

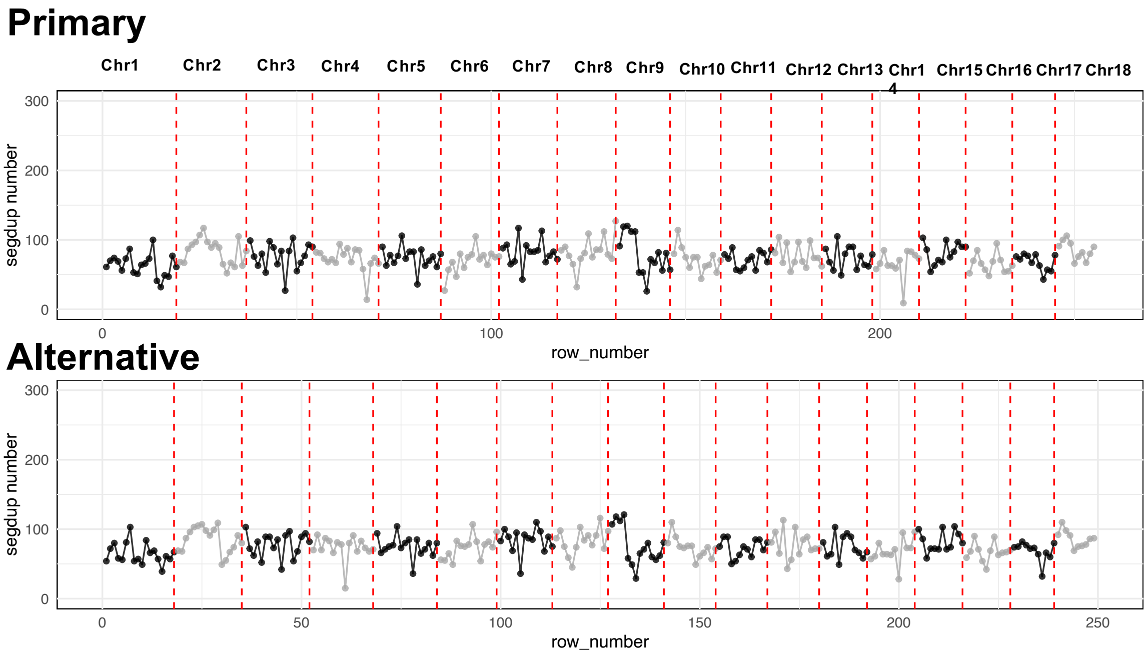


**Figure S8.** Counts of segmental duplication genes in the primary and alternative assemblies using a 5 mega-base pair sliding window.

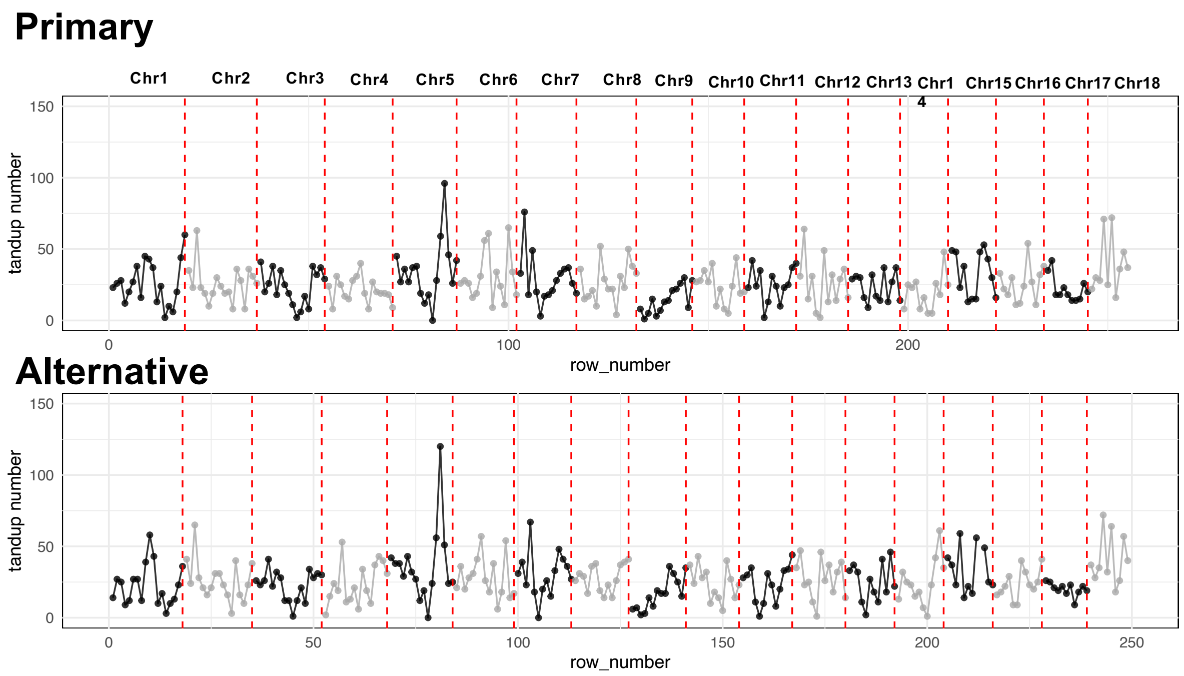


**Figure S9.** Counts of tandem duplication genes in the primary and alternative assemblies using a 5 mega-base pair sliding window.

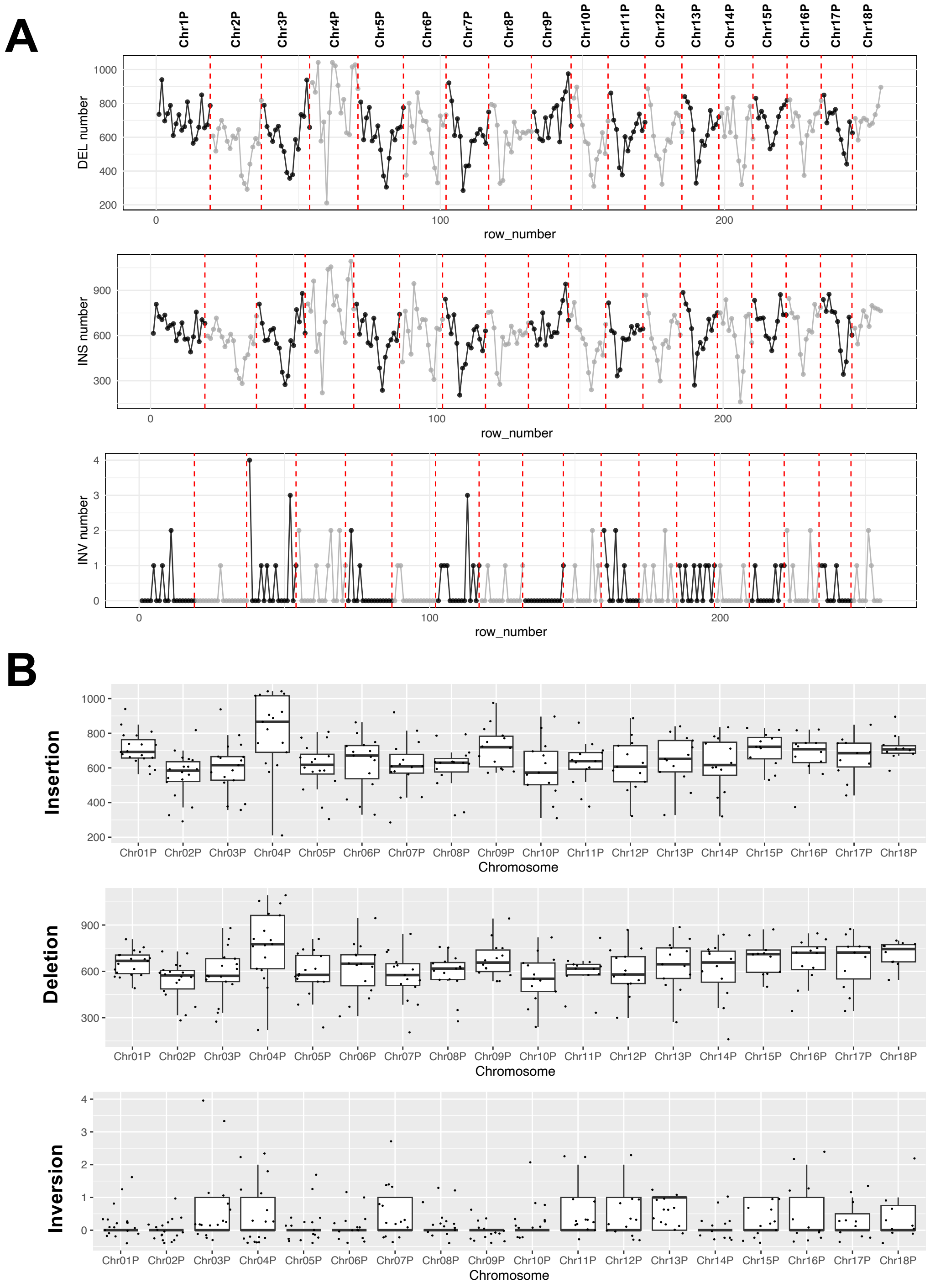


**Figure S10. A.** Counts of structural variations (insertion, deletion,and inversion)  in the primary assembly using a 5 mega-base pair sliding window. **B.** Boxplots for the counts of of structural variations using a 5 mega-base pair sliding window.

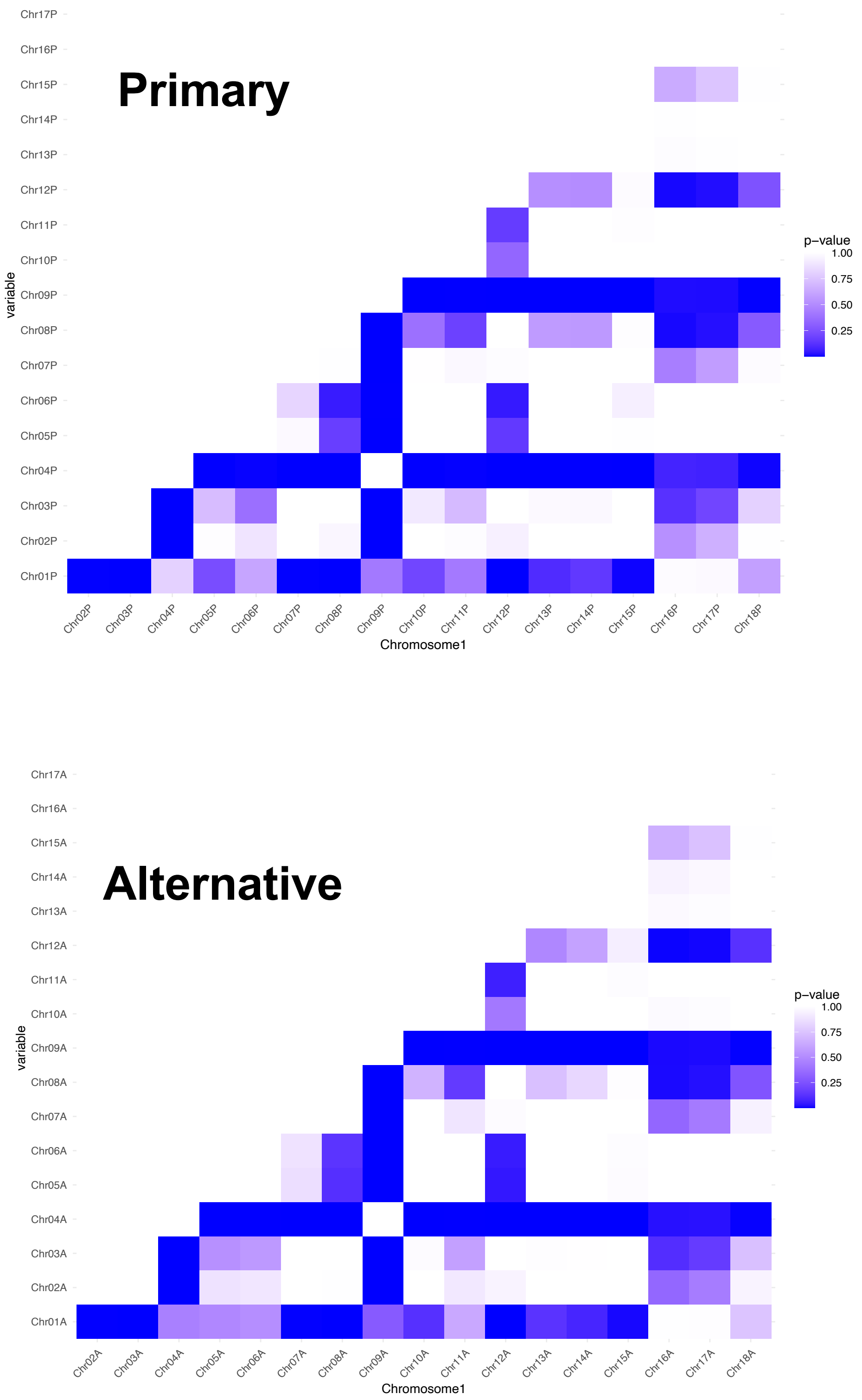


**Figure S11.** Heatmap of *p*-values obtained from Tukey's post-hoc Honestly Significant Difference (HSD) test for BUSCO counts. The color gradient represents the significance levels of pairwise comparisons.

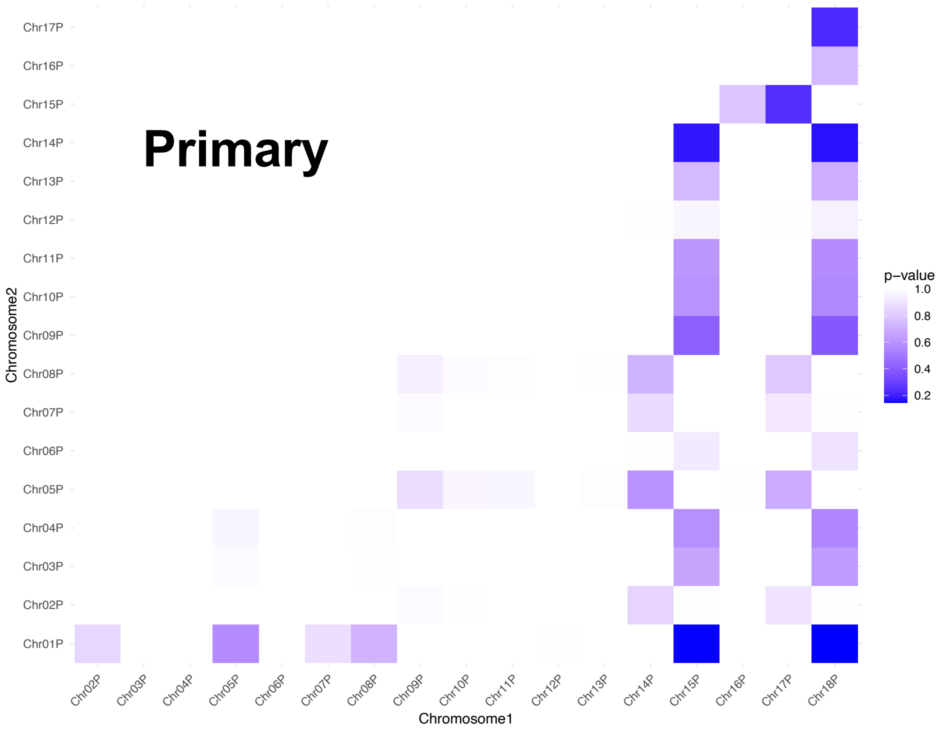


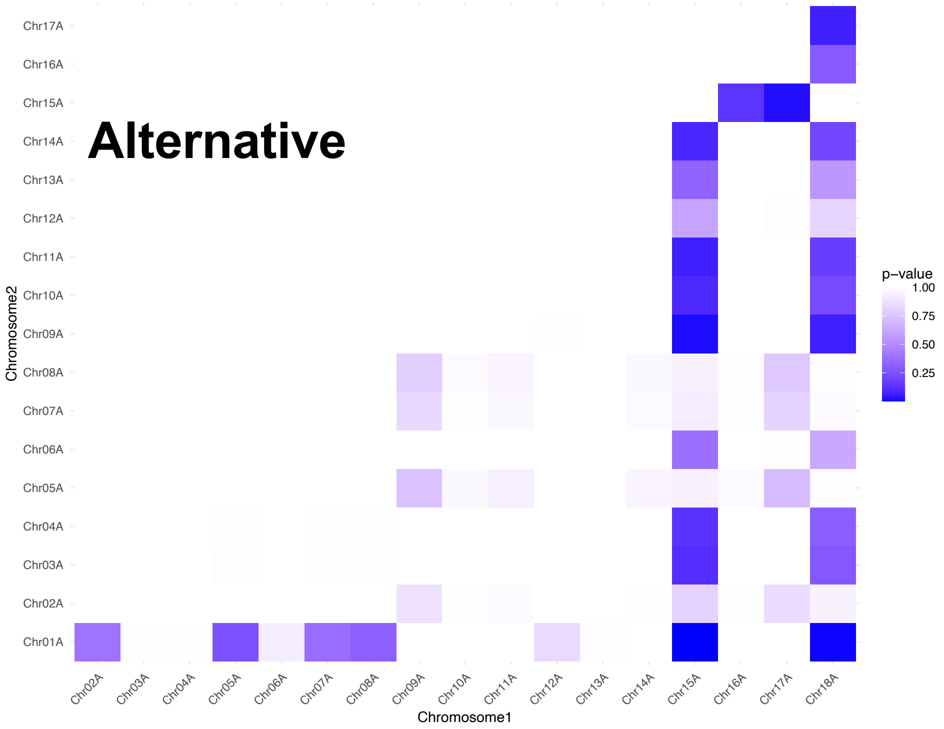


**Figure S12.** Heatmap of *p*-values obtained from Tukey's post-hoc Honestly Significant Difference (HSD) test for gene counts. The color gradient represents the significance levels of pairwise comparisons.

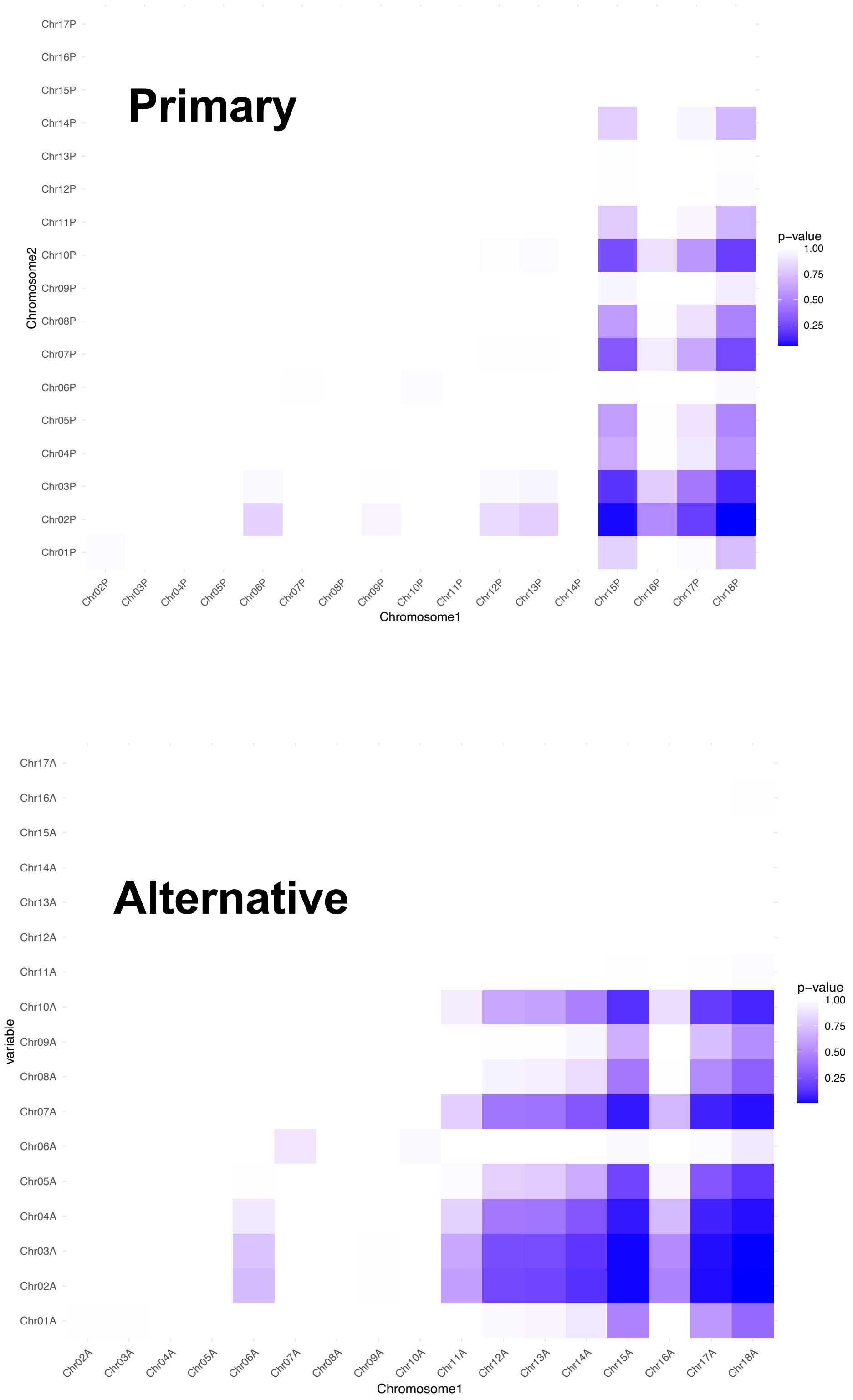


**Figure S13.** Heatmap of *p*-values obtained from Tukey's post-hoc Honestly Significant Difference (HSD) test for repeat counts. The color gradient represents the significance levels of pairwise comparisons.

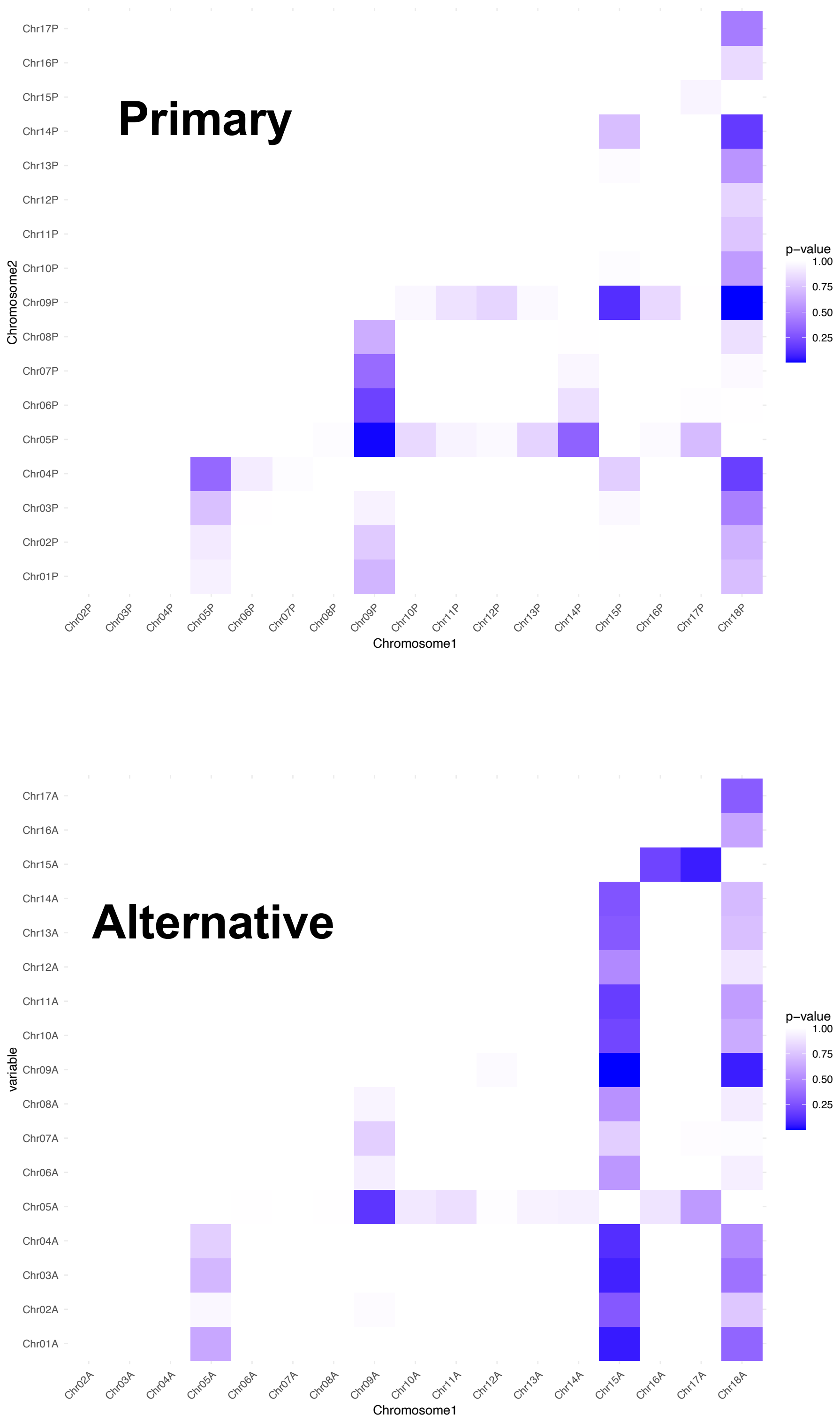


**Figure S14.** Heatmap of *p*-values obtained from Tukey's post-hoc Honestly Significant Difference (HSD) test for segmental duplication counts. The color gradient represents the significance levels of pairwise comparisons.

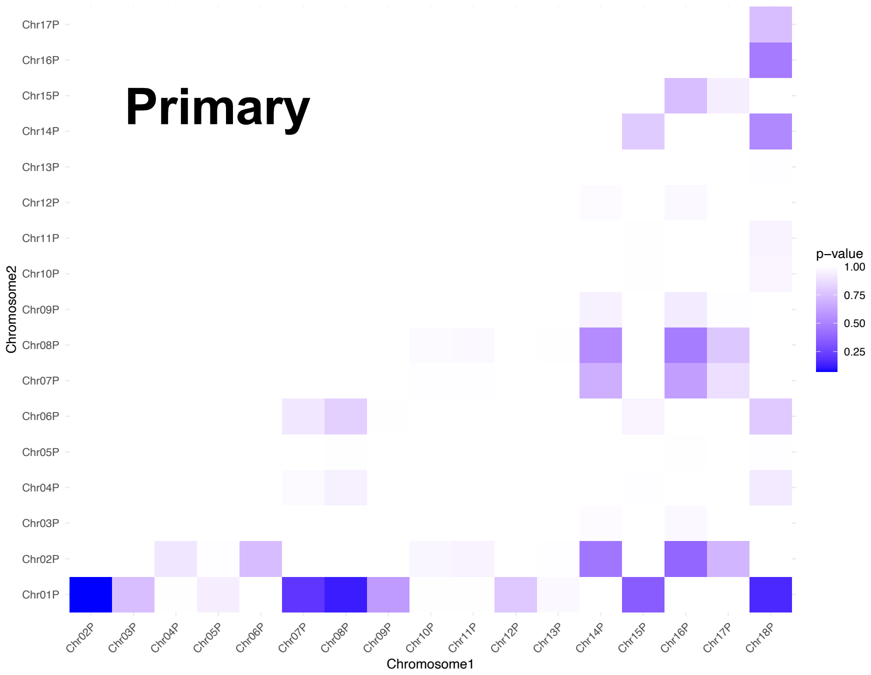


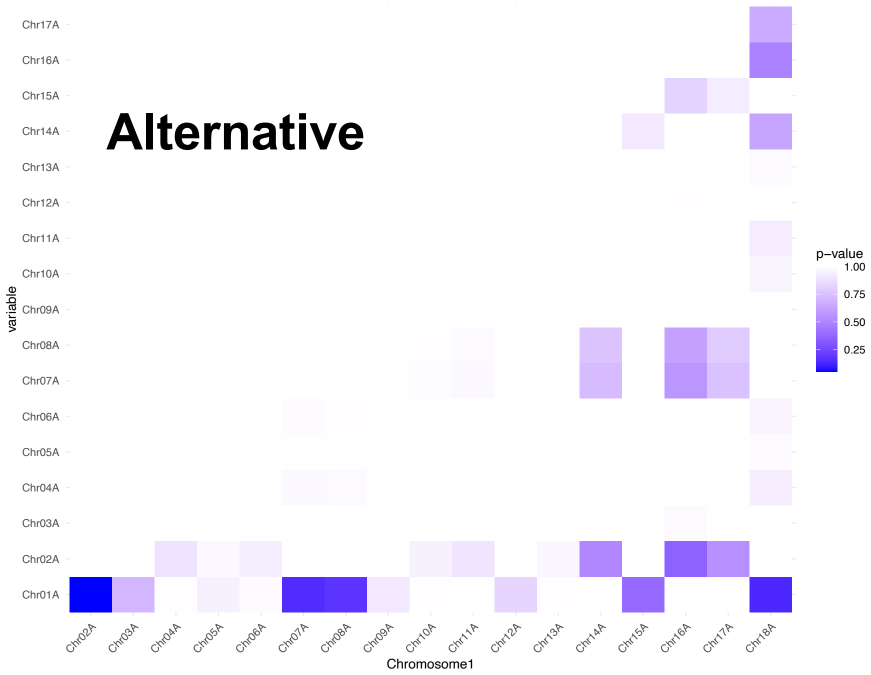


**Figure S15.** Heatmap of *p*-values obtained from Tukey's post-hoc Honestly Significant Difference (HSD) test for tandem duplication counts. The color gradient represents the significance levels of pairwise comparisons.

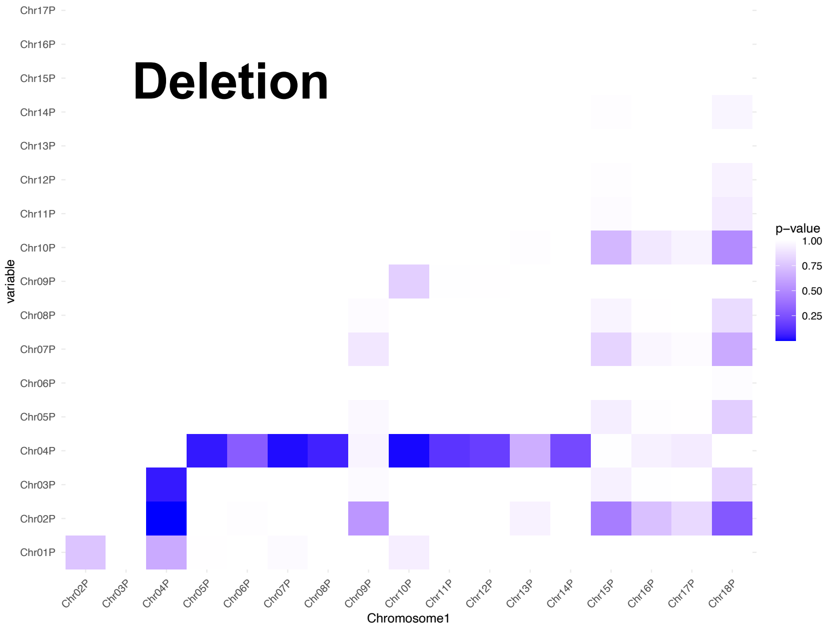


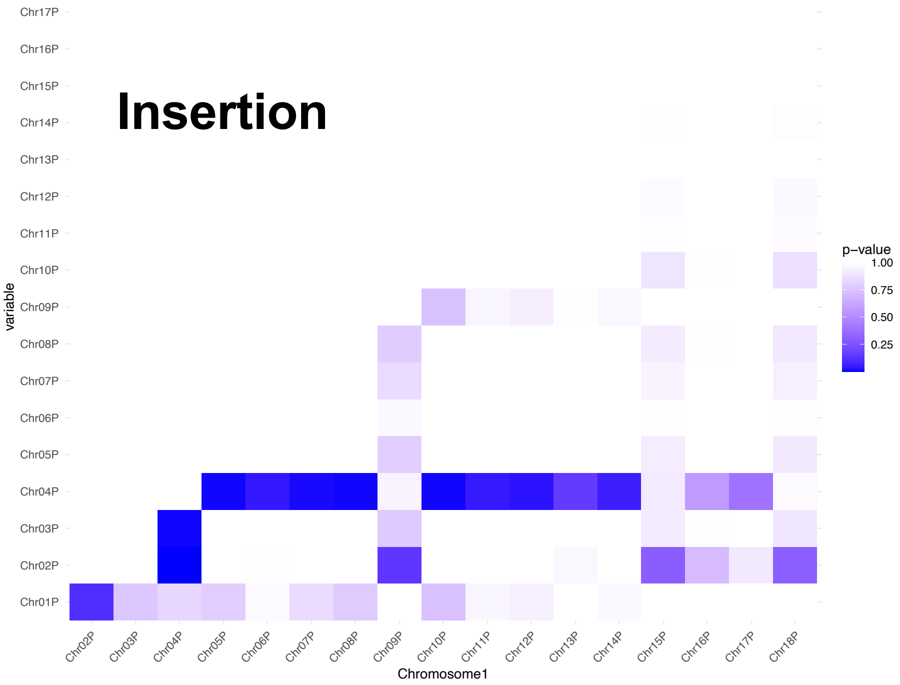


**Figure S16.** Heatmap of *p*-values obtained from Tukey's post-hoc Honestly Significant Difference (HSD) test for deletion and insertion counts. The color gradient represents the significance levels of pairwise comparisons.

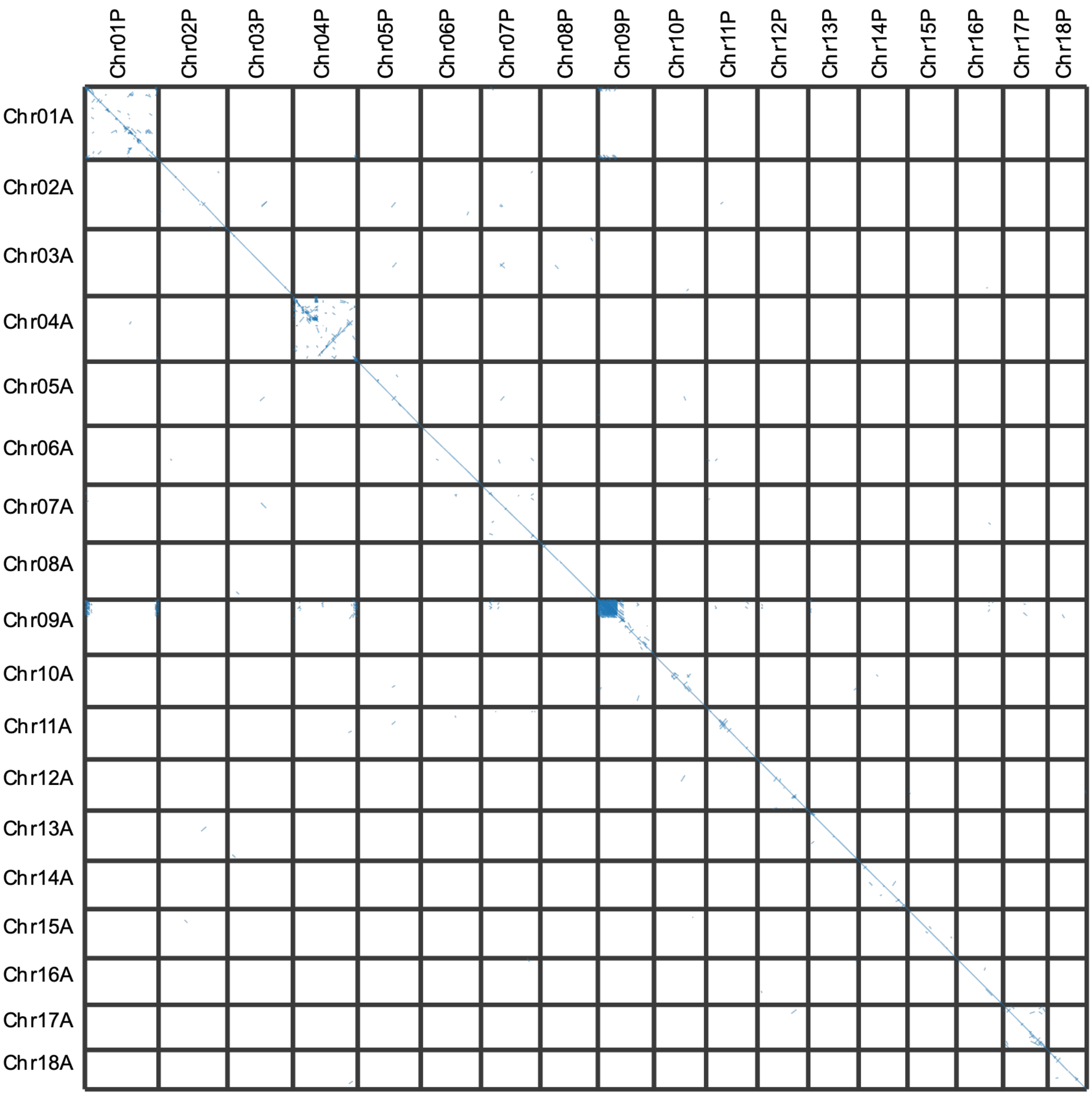


**Figure S17.** Interchomosomal and intrachomosomal syntenic gene blocks (primary vs alternative assemblies).

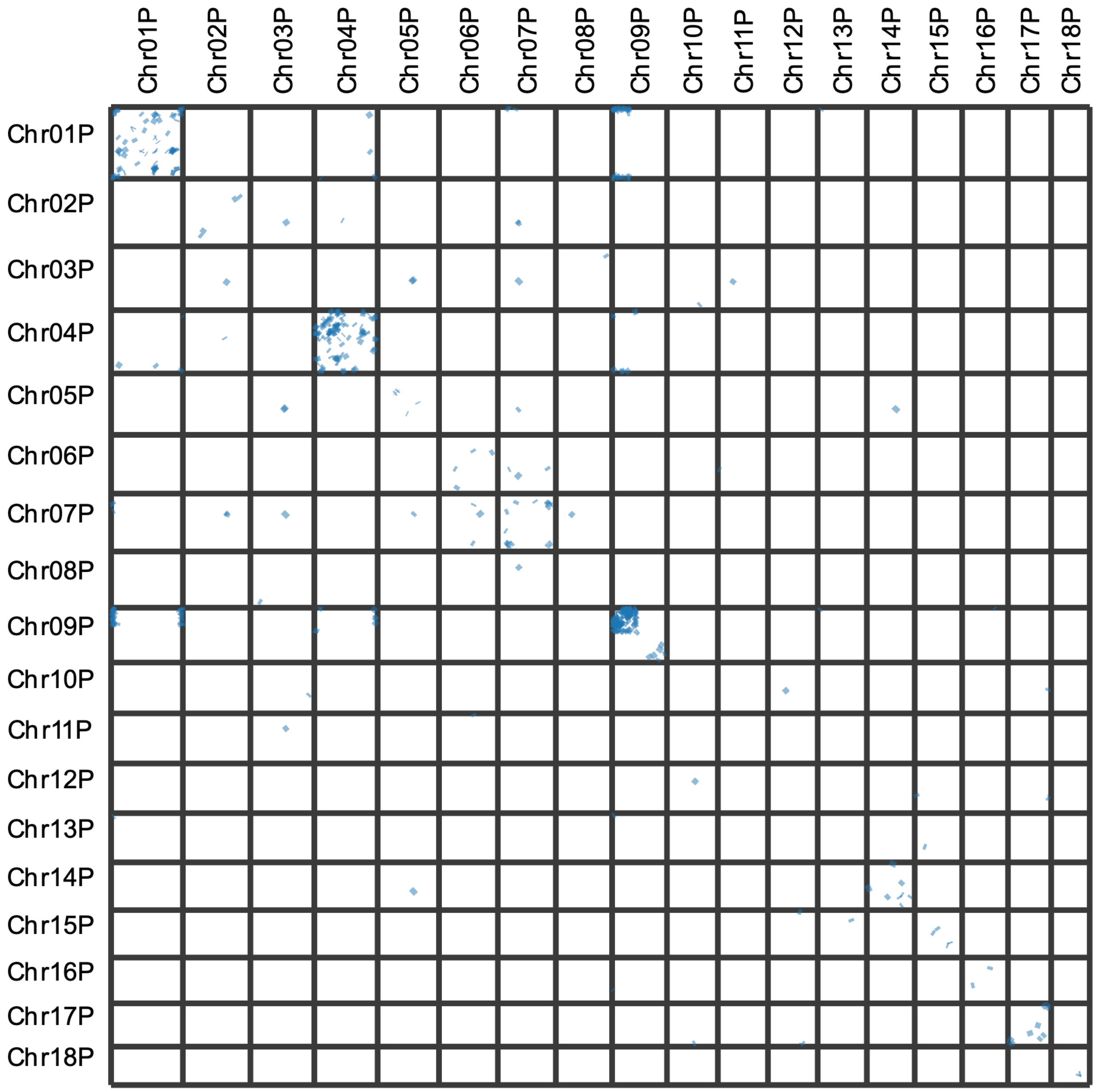


**Figure S18.** Interchomosomal and intrachomosomal syntenic gene blocks (primary vs primary assemblies).

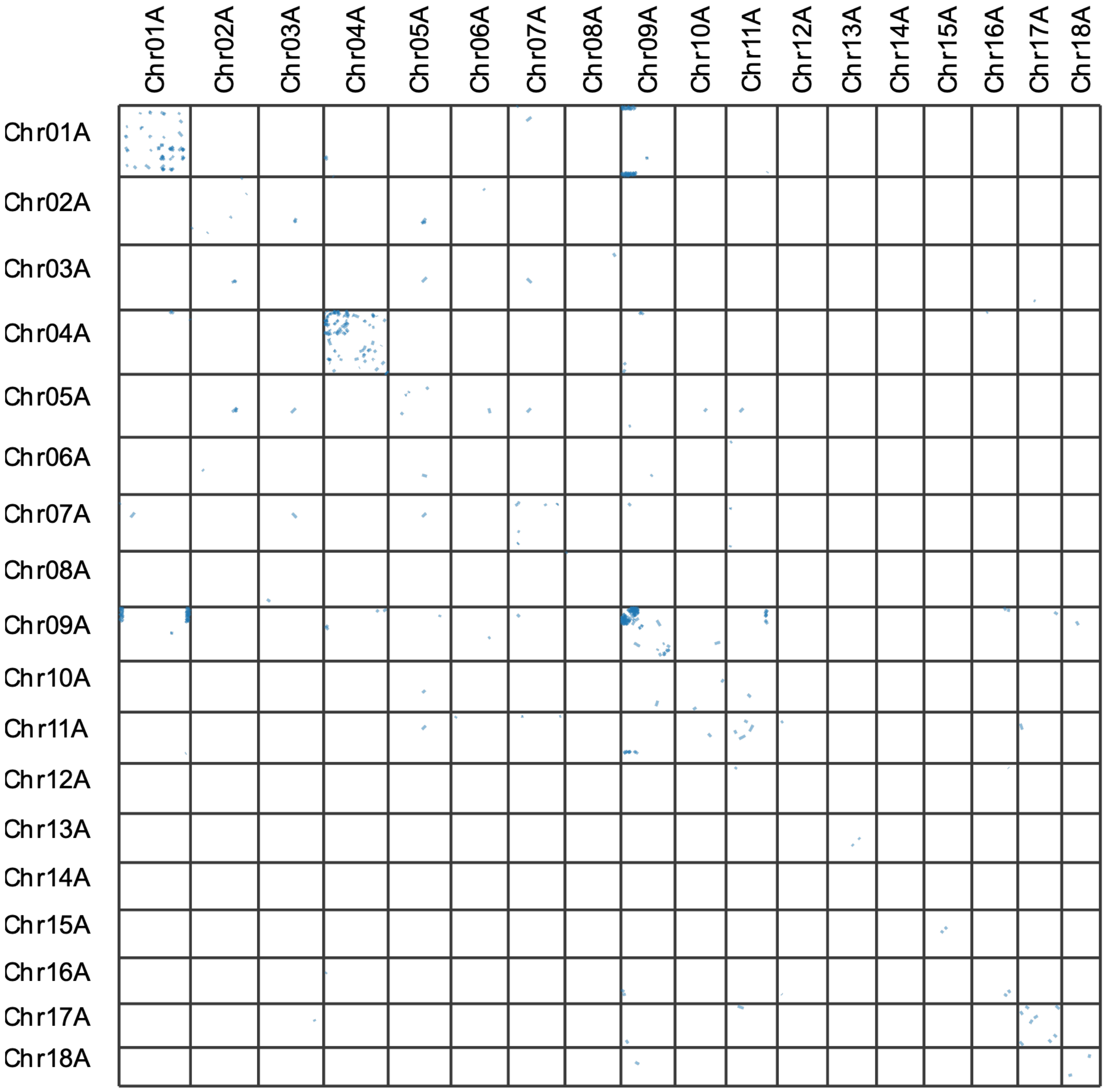


**Figure S19.** Interchomosomal and intrachomosomal syntenic gene blocks (alternative vs alternative assemblies).

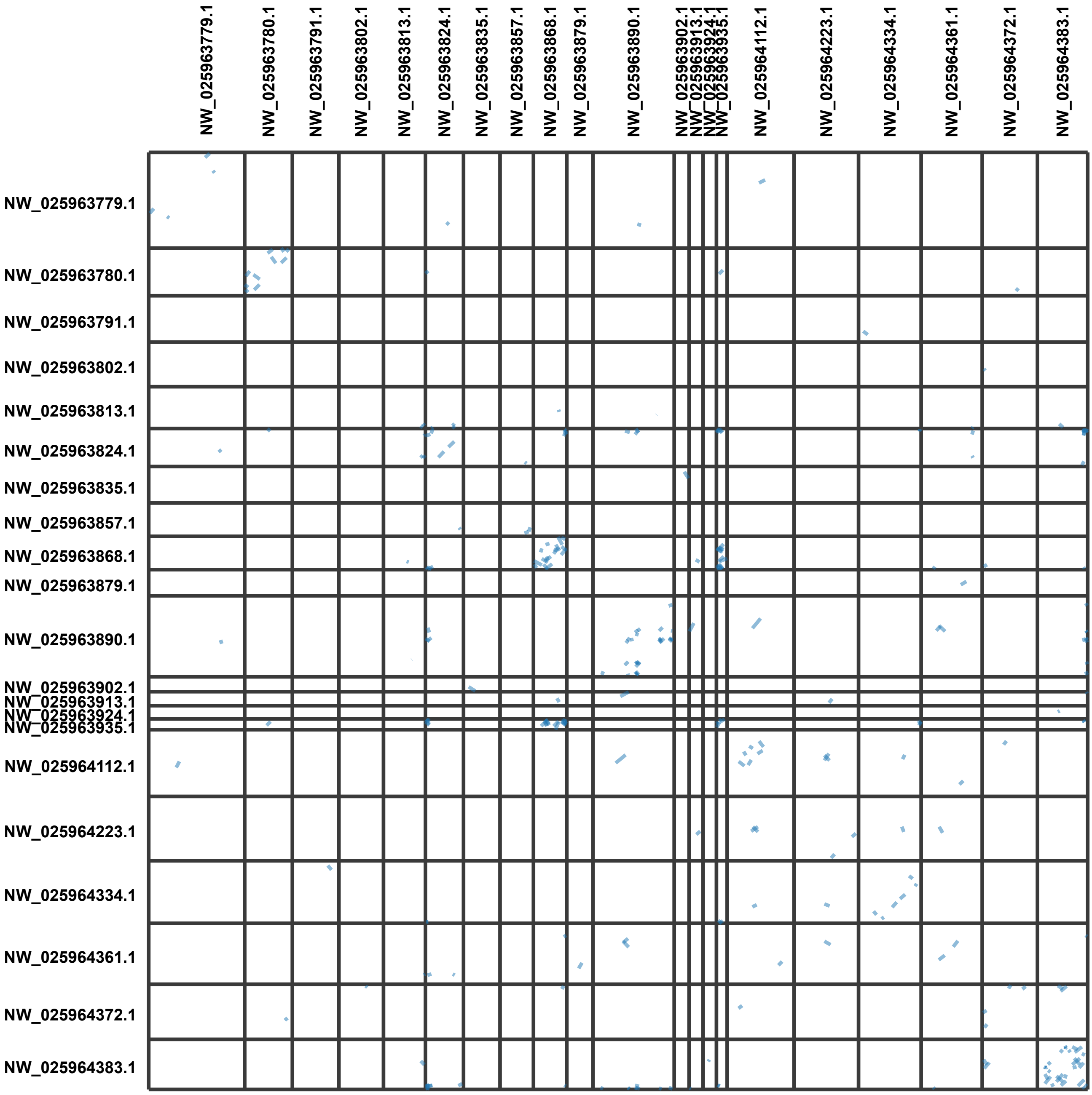


**Figure S20.** Interchomosomal and intrachomosomal syntenic gene blocks within *Haliotis rufescens* scaffolds.

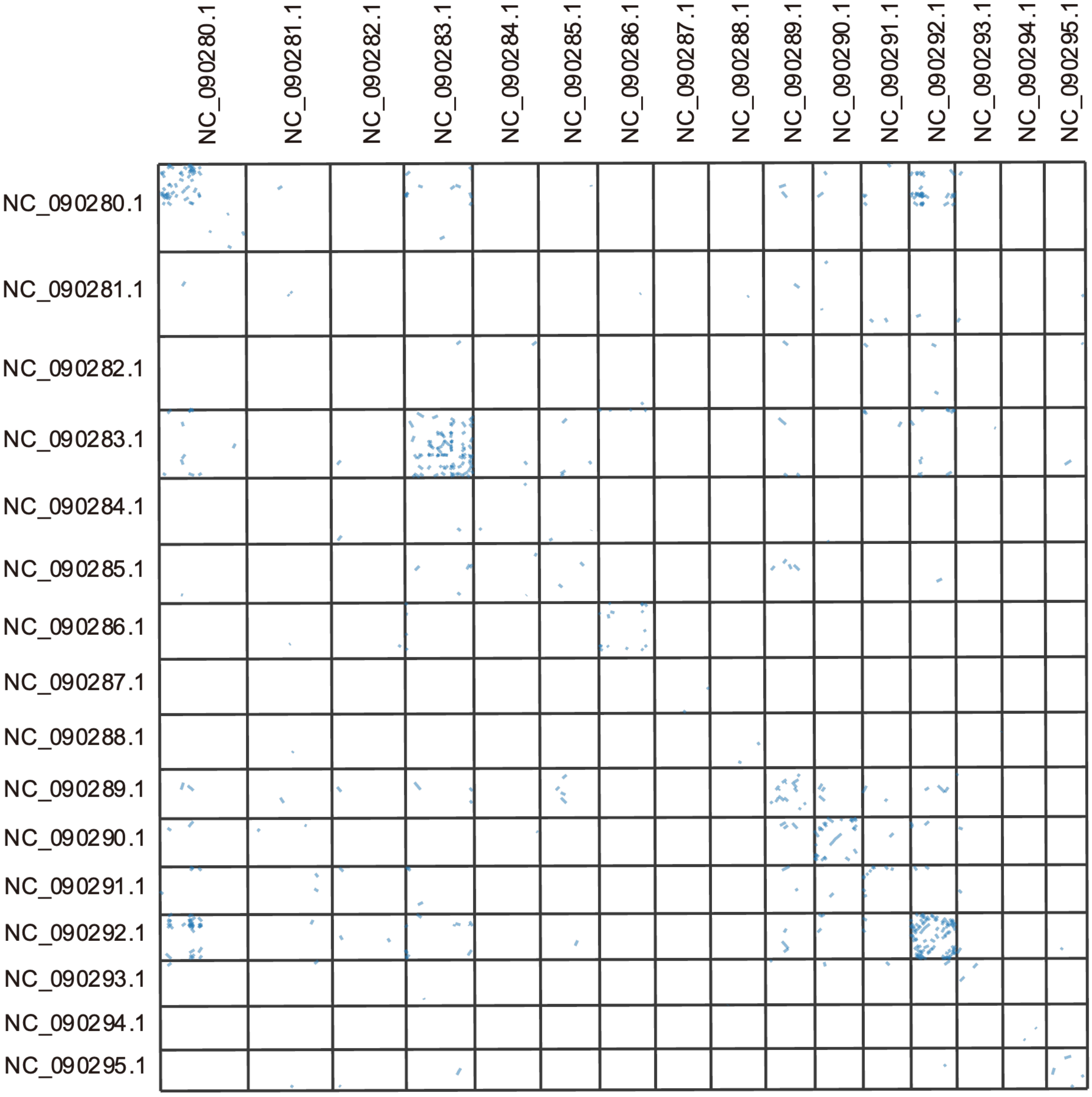


**Figure S21.** Interchomosomal and intrachomosomal syntenic gene blocks within *Haliotis asinina* scaffolds.

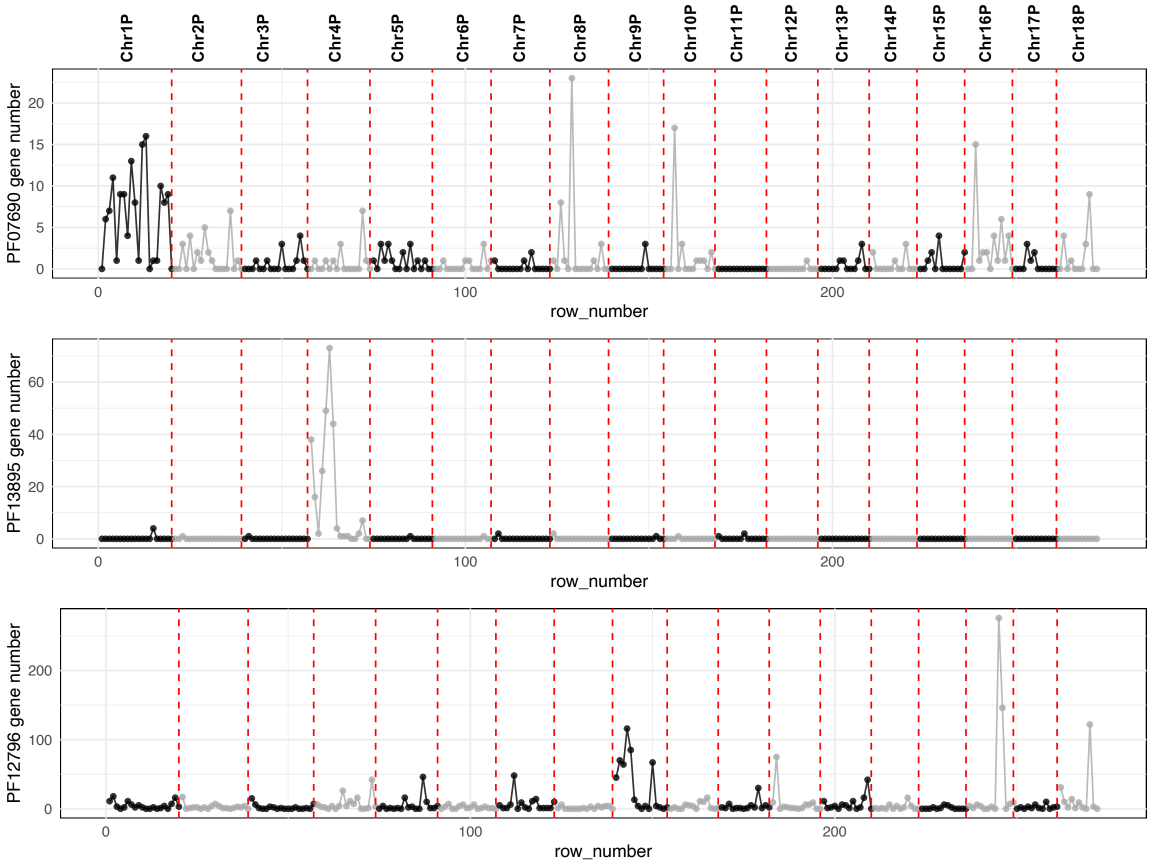


**Figure S22.** Counts of genes with each of three enriched Pfam domain (PF07690 in Chr1, PF13895 in Chr4, and PF12796 in Chr9) in the primary assembly using a 5 mega-base pair sliding window.

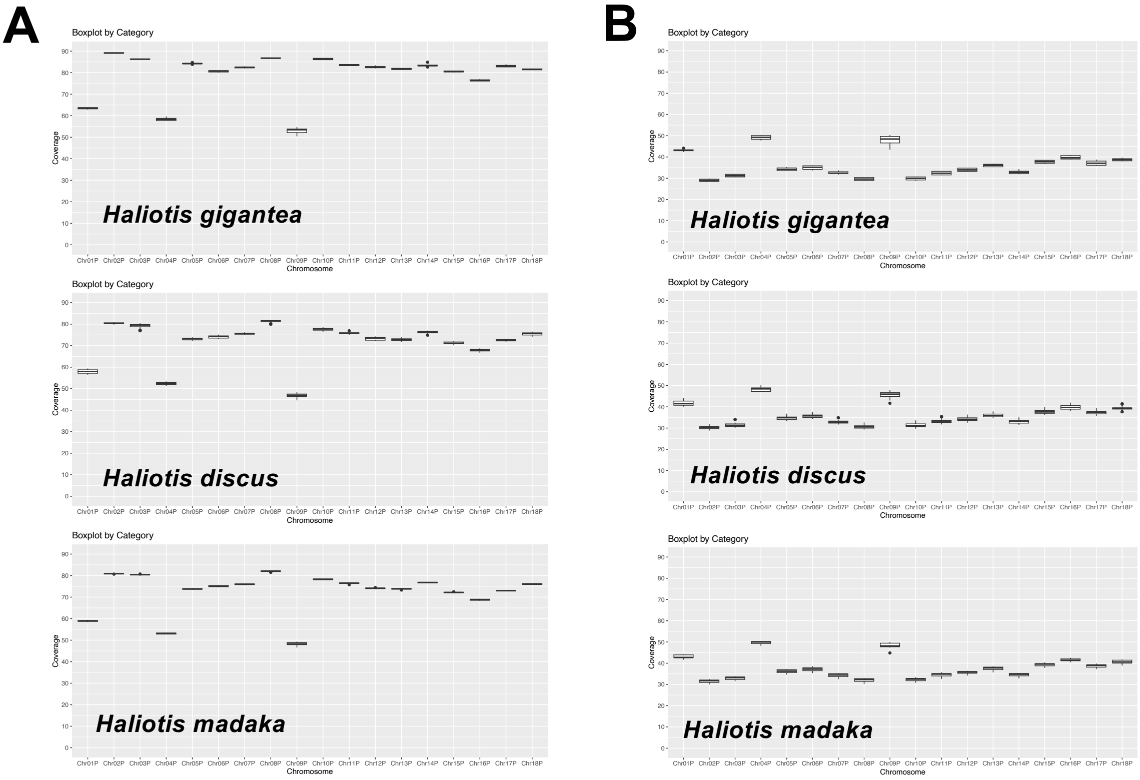


**Figure S23.** Percentage of genomic regions where short reads mapped uniquely (A) and multiply (B) on each chromosome.

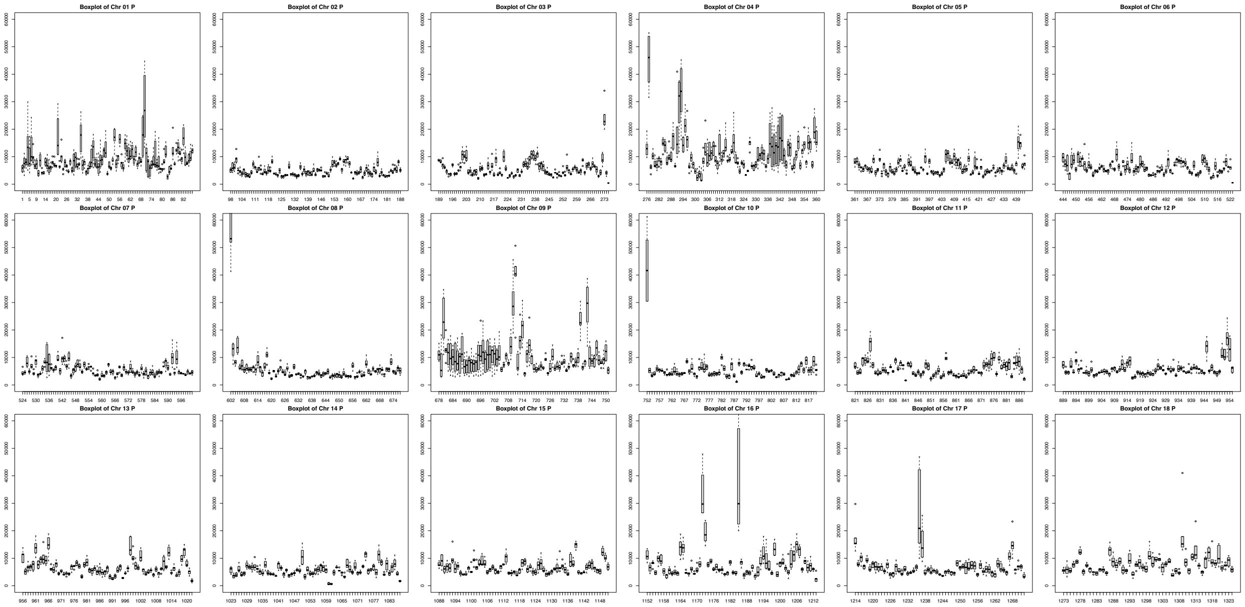

**Figure S24.** Normalized numbers of mapped reads from *Haliotis gigantea* individuals in each chromosome of primary assembly.

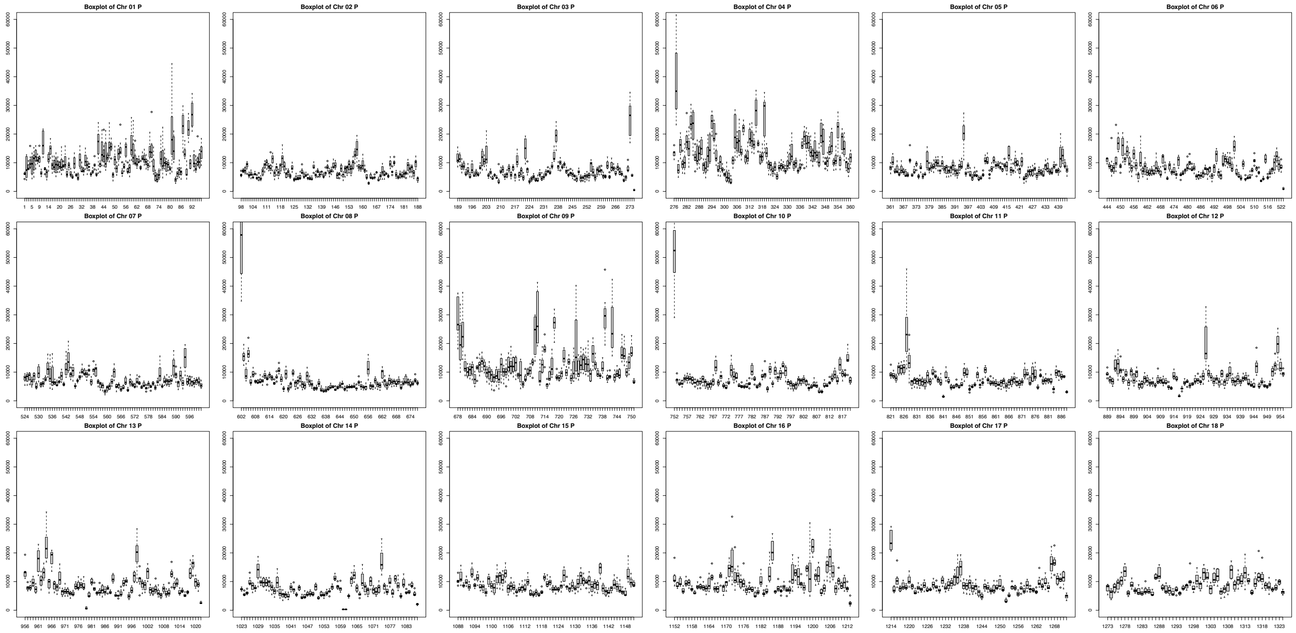


**Figure S25.** Normalized numbers of mapped reads from *Haliotis madaka* individuals in each chromosome of primary assembly.

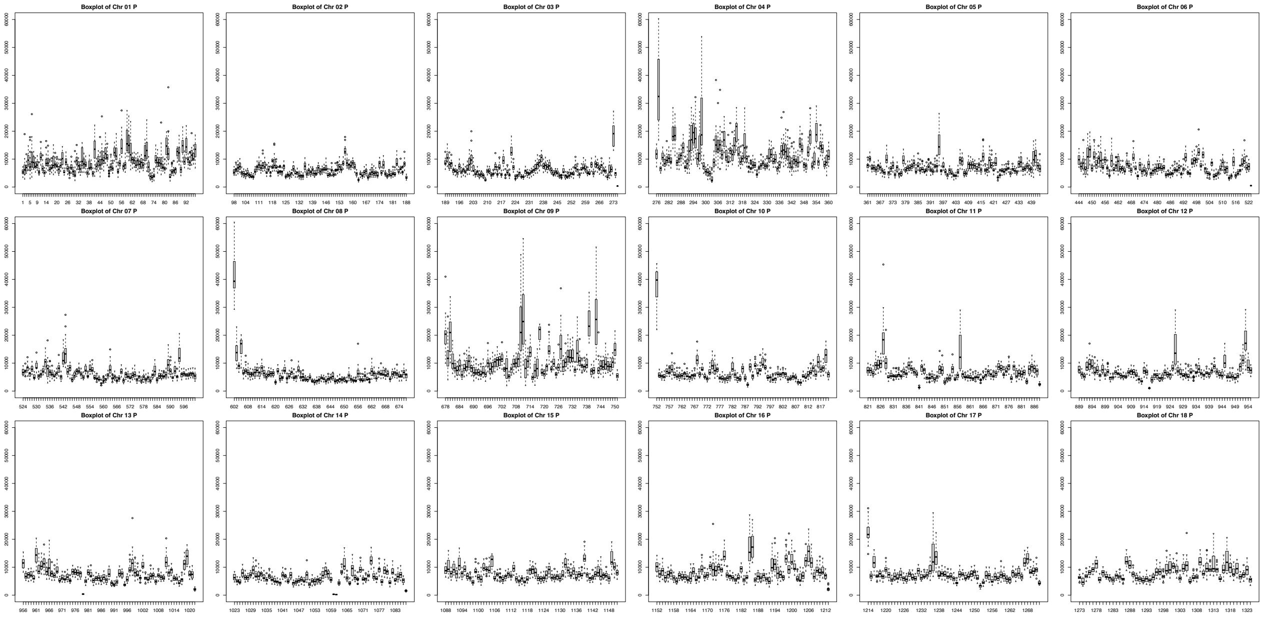


**Figure S26.** Normalized numbers of mapped reads from *Haliotis discus* individuals in each chromosome of primary assembly.
